# Extending structural surfaceomics to identify aberrant conformations of tumor surface proteins as potential immunotherapy targets

**DOI:** 10.64898/2026.05.15.721813

**Authors:** Audrey Kishishita, Sabine Cismoski, Tianna Grant, Rucha Deo, Sanjana Prudhvi, Catherine Sue, Abhilash Barpanda, Clinton Yu, Sanjyot Shenoy, Sarah Berman, Audrey G. Reeves, Haolong Li, Tianyi Liu, Akul Naik, Deeptarup Biswas, Fenglong Jiao, Yi He, Matthew Hancock, Radhika Dalal, Arther Zalevsky, Michael R. Hoopmann, Chun Jimmie Ye, Rosa Viner, Felix Feng, Kamal Mandal, Robert L. Moritz, Ignacia Echeverria Riesco, Andrej Sali, James A. Wells, Sanjeeva Srivastava, Lan Huang, Arun P. Wiita

## Abstract

The complement of tumor cell surface proteins, or “surfaceome”, is a rich source of potential immunotherapy targets. To move beyond expression-based target discovery, we previously described “structural surfaceomics,” combining crosslinking mass spectrometry (XL-MS) with surface protein biotinylation to identify conformation-selective targets. In our prior work, we applied this method to a single model of acute myeloid leukemia (AML), identifying active integrin beta-2 as a promising target. Here, we expand structural surfaceomics to identify additional immunotherapy targets and surface protein biology across additional models of AML, multiple myeloma, and prostate cancer, as well as donor peripheral blood mononuclear cells. Utilizing these models and different chemical crosslinkers, we compile an extensive database of 5,209 crosslinks. We characterize both shared and unique crosslink-based features, identifying 1,612 disease model-specific crosslinks, including 212 potentially defining tumor-specific conformations based on distance constraint violations relative to AlphaFold predictions. We further implement a suite of emerging modeling tools to predict tumor-specific protein structures. We probe crosslinking patterns suggesting multiple myeloma-specific CD48 and AML-specific integrin α1/β4 heterodimer conformations. This work establishes a resource for cancer structural biology by implementation of structural surfaceomics. Our findings also point toward more realistic protein design models, potentially enabling systematic detection of targetable cancer-specific epitopes for next-generation immunotherapies.

## INTRODUCTION

The landscape of surface proteins on cancer cells, or the cancer “surfaceome”, governs tumor interactions with its microenvironment and represents a primary interface for therapeutic intervention^1–3^. Tumor surface antigen–targeted immunotherapies, including monoclonal antibodies (mAbs), antibody–drug conjugates (ADCs), bispecific T cell engagers (TCEs), and chimeric antigen receptor (CAR) T cells, have become the most exciting classes of cancer therapeutics^4^. Prominent examples include widely prescribed therapeutics such as cetuximab, a mAb targeting the surface antigen EGFR in colorectal cancer^5–7^, and tisagenlecleucel, a CAR T-cell targeting the surface antigen CD19 in B-cell leukemia^8,9^. However, most surface antigens are shared between tumor and normal tissues, leading to “on-target, off-tumor” (OTOT) toxicities that limit therapeutic index. Such toxicities can range from clinically tolerable phenotypes to fatal organ damage, underscoring the narrow therapeutic index of many surface targets^10,11^. Thus, identifying tumor-selective surface antigens remains a central challenge in cancer immunotherapy.

To date, the vast majority of target discovery efforts in the cancer immunotherapy field have focused on using genomics and transcriptomics datasets to identify surface proteins that may be upregulated in tumors versus normal tissues^12,13^. However, these approaches prioritize transcript abundance, whereas antigen-specific immunotherapies fundamentally rely on structural epitopes for target recognition, typically via an antibody or antibody fragment. Tumor-specific conformations may therefore create selective epitopes even when protein expression is unchanged.

To address this challenge, we recently developed “structural surfaceomics,” leveraging the hypothesis that 3-dimensional conformations of tumor surface proteins may present novel immunotherapy targets with distinct, tumor-specific epitopes independent of expression level^14–16^. Many plasma membrane proteins adopt distinct conformational states in response to, and thus may be driven by aberrant tumor signaling, metabolism, cell-cell interactions, or other biological processes^17^. In our initial study, we combined cell-surface biotinylation with crosslinking mass spectrometry (XL-MS) to identify conformation-selective targets and validated an active form of integrin β2 as a tumor-selective CAR T target in acute myeloid leukemia (AML)^18^. Notably, this epitope would not have emerged from conventional abundance-based discovery approaches, as integrin β2 is broadly expressed on normal hematopoietic cells in the closed, inactive conformation.

While highly promising, our initial approach was limited in its scope. Analyses were restricted to a single AML model, and structural interpretation relied on manually mapping crosslinks onto available Protein Data Bank (PDB) structures, which is constrained by the under-representation of membrane proteins. Recent advances in protein structure prediction, particularly AlphaFold, now enable systematic modeling of membrane protein architectures and multimers, creating an opportunity to computationally evaluate crosslink-defined conformations across the surface proteome^19–21^.

Here, we optimize and scale structural surfaceomics to systematically interrogate conformational heterogeneity across multiple cancer models and matched non-malignant cells. By integrating orthogonal crosslinking strategies with recently-developed structure-guided computational modeling ^22^, we identify disease-associated conformations that nominate previously unrecognized immunotherapy targets, including a multiple myeloma–specific conformation of CD48 and an AML-specific heterodimeric integrin α1/β4 complex, followed by emerging deep learning-based tools to model physically realistic cancer-specific protein conformations from experimental constraints^23–27^. Beyond individual proteins, our analysis suggests that malignant transformation may remodel higher-order membrane protein assemblies, revealing an additional layer of surfaceome organization. This work arrives at a critical convergence point: enhanced LC-MS/MS sensitivity, maturation of XL-MS techniques and data processing^28–31^, the emergence of AI-driven structural prediction tools^23,24,32,33^, and the urgent clinical need for new immunotherapy targets^34–36^. Furthermore, our dataset of >5,000 crosslinked peptides from the surface-enriched proteome represents a unique resource for studying the structural biology of membrane proteins in cancer. Together, these findings establish conformational heterogeneity as a systematic and targetable feature of the cancer surfaceome and provide a generalizable framework for epitope-centric target discovery.

## Results

### Optimization of surface biotinylation and crosslinking workflow

To systematically identify tumor-specific protein conformations at the cell surface, we previously developed structural surfaceomics, integrating cell surface biotinylation with crosslinking mass spectrometry (XL-MS) (**Fig. 1A**). Live cells undergo crosslinking with either disuccinimidyl sulfoxide (DSSO), a homobifunctional cleavable NHS-ester crosslinker that reacts with primary amines on lysine residues, or PhoX (phosphonate-tagged crosslinker), an IMAC-enrichable NHS-ester crosslinker with a shorter, less flexible spacer arm (molecular structures shown in **Fig. 1A**). While both crosslinkers react with lysine residues, DSSO’s cleavable sulfoxide linkage facilitates identification of crosslinked peptide pairs via MS^3^ fragmentation, whereas PhoX’s phosphonate moiety enables selective IMAC-based enrichment prior to LC-MS/MS analysis, followed by surface protein biotinylation. This orthogonal labeling strategy—lysine-targeting crosslinkers and glycan-targeting biotinylation—avoids competition for labeling sites while maintaining cell viability (**Fig. 1B**). To validate selective surface protein enrichment, we selected CD48 as a positive control due to its known high expression in multiple myeloma relative to peripheral blood mononuclear cells (PBMCs)^37–40^. Immunoprecipitation followed by Western blot confirmed specific enrichment of high molecular weight biotinylated CD48 in crosslinked myeloma samples, consistent with successful crosslink formation^41^ (**Fig.1C**).

**Figure 1.**
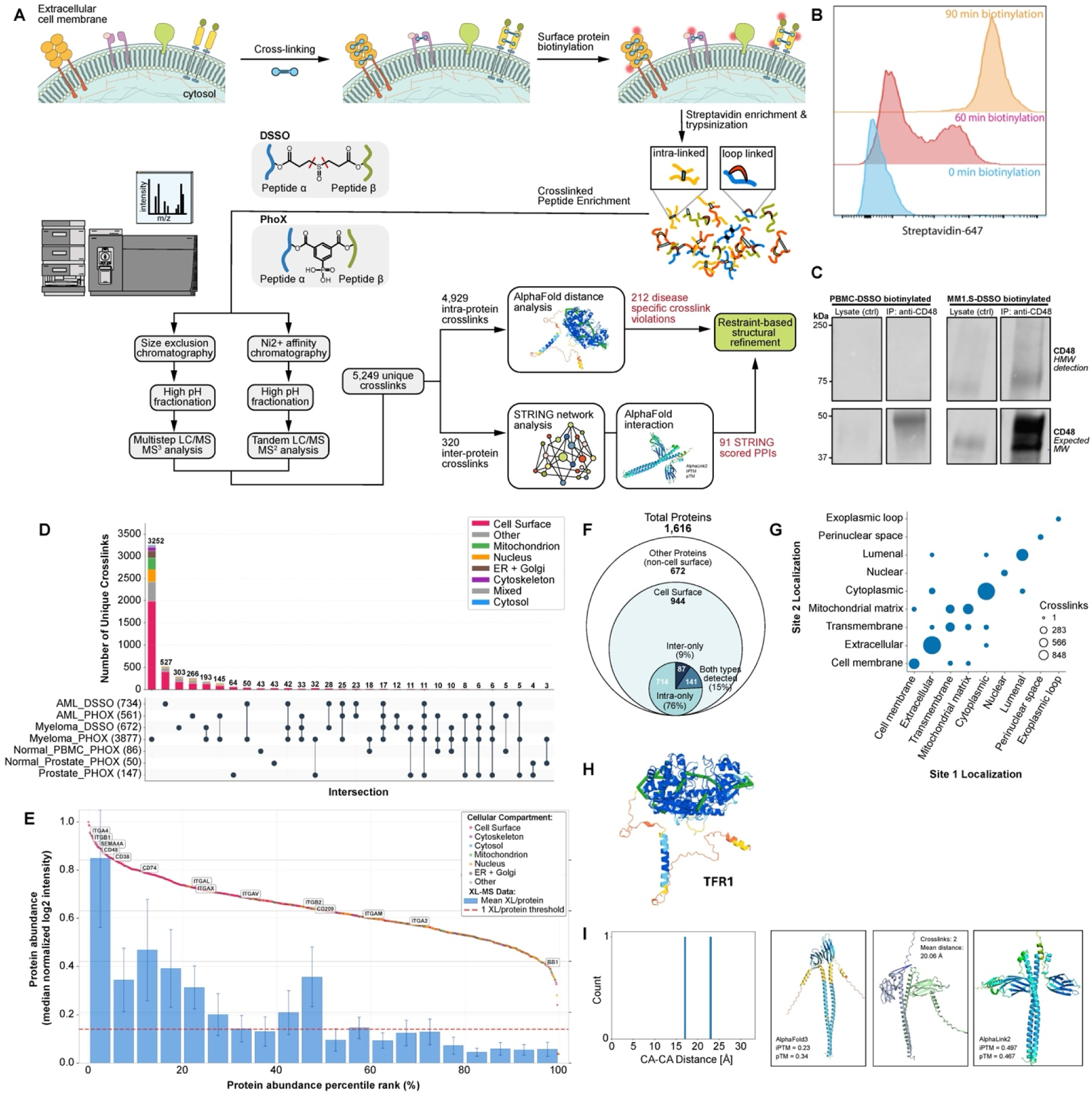
Structural Surfaceomics Workflow and Dataset Overview. **(A)** Structural surfaceomics workflow: live cells undergo surface biotinylation followed by crosslinking with DSSO or PhoX, streptavidin affinity enrichment, and trypsinization. Crosslinked peptides are separated by 2D fractionation (SEC or IMAC, then high-pH RP) and analyzed by LC-MS/MS, with data processed through pLink, AlphaCross-XL, and MODELLER/AlphaLink pipelines. **(B)** Flow cytometry confirms 90-minute biocytin hydrazide incubation achieves optimal surface labeling. **(C)** Western blot of anti-CD48 immunoprecipitation from PBMC-DSSO and MM1.S-DSSO lysates confirms specific enrichment of surface-biotinylated, high-molecular-weight CD48 in myeloma cells following the structural surfaceomics workflow. **(D)** UpSet plot of unique crosslinks across 9 cancer cell line datasets, color-coded by subcellular compartment localization. **(E)** Protein abundance rank plot with mean crosslinks per protein per percentile bin (blue bars); expression values for all proteins from bottom-up surfaceomics analysis of the malignancy models (rank plot), where high-abundance proteins show greater XL-MS detection. **(F)** Of 1,616 proteins identified, 944 (58%) are cell surface-annotated; surface proteins are classified by crosslink type: intra-only (76%), intra- and inter-protein (15%), and inter-only (9%). **(G)** Topology-based crosslink classification confirms surface-selective enrichment. **(H–I)** AlphaCross-XL pipeline maps intra-protein crosslinks to AlphaFold2 monomers **(H)** and inter-protein crosslinks to AlphaFold-Multimer predictions **(I)**, enabling proteome-scale distance constraint validation and conformational analysis. Average cross-link per protein abundance percentile. iBAQ values for all proteins found in a bottom-up analysis of *N. meningitidis* (gray) with proteins found cross-linked in an in vivo experiment (blue). The bar plots represent the average cross-link per protein calculated per 5% protein abundance percentile with a cut off at 1 cross-link per protein, highlighted as a dashed red line.

To maximize structural surfaceome coverage, we compared two glycan-dependent biotinylation strategies. Flow cytometry confirmed that 90-minute biocytin hydrazide incubation achieved complete surface labeling (**Fig. 1B**). We then compared biocytin hydrazide to an enzymatic alternative—wheat germ agglutinin-horseradish peroxidase (WGA-HRP)—which biotinylates N-glycans and sialic acid residues with shorter reaction times^42,43^. Testing both approaches on DSSO-crosslinked suspension cells across blood cancers revealed that biocytin hydrazide yielded 1.65-fold more crosslinks (**Fig. S1**), while both methods identified similar numbers of surface proteins (111 vs 135 proteins, **Fig. S2**). Following these results, we selected biocytin hydrazide as the surface protein enrichment strategy for all subsequent datasets in this study for consistency with previous results.

To enhance crosslink detection, we employed two-dimensional fractionation tailored to each crosslinker chemistry^44,45^. For DSSO samples, we used size exclusion chromatography (SEC) followed by high pH reversed-phase (HpH-RP) fractionation^46^. For PhoX samples, we leveraged the phosphonate tag for immobilized metal affinity chromatography (IMAC) as a first dimension of enrichment^47^. HpH-RP as the second dimension after IMAC yielded 5.8-fold more detected crosslinks than SEC in PhoX-labeled AML samples (**Fig. S3**), establishing IMAC-HpH-RP as the optimal workflow for these datasets.

### Comprehensive structural surfaceomics across cancer models

To extend the results from our prior study^18^, we applied structural surfaceomics to six cancer cell lines models representing AML, myeloma, and prostate cancer (AML: THP-1, MV4-11, NOMO-1; multiple myeloma: AMO-1, MM.1S; prostate cancer: 22Rv1) and two non-malignant comparators (donor peripheral blood mononuclear cells (PBMCs), PNT2 immortalized benign prostatic epithelium cell line). Across these 11 datasets, we identified 5,209 unique crosslinks from 19,907 crosslinked spectral matches after quality filtering (**Fig. 1D** and **Methods**). Unsupervised clustering of high-confidence crosslinks grouped samples by disease type, with myeloma samples showing the strongest within-disease concordance (mean Jaccard similarity = 0.50). Notably, two independent myeloma cell lines (AMO1 and MM.1S) shared 87% of their high-abundance crosslink profiles (**Fig. 1D**), confirming that crosslinking patterns capture disease-associated structural features.

In the overall dataset, crosslink detection correlated with protein abundance, with surface proteins showing highest crosslink density (**Fig. 1E** and **Supplementary Data 1**); this finding of greater resolution for high abundance proteins is typical for mass spectrometry-based approaches. Of 1,616 proteins identified, 944 (58%) were cell surface-annotated (**Fig. 1F**), confirming surfaceome enrichment. Topological analysis revealed that 48.3% of crosslinked residues (n=1,977) mapped to cell surface domains (extracellular, transmembrane, and membrane-associated regions) (**Fig. 1G**), consistent with surface-selective biotinylation. The remaining crosslinks included cytoplasmic (38.6%, n=1,582), lumenal (7.4%, n=302), and mitochondrial matrix (3.5%, n=142) localizations. Next, employing our recently-described AlphaCross-XL software^22^—a proteome-scale pipeline for automated mapping of crosslinked peptides onto 3D protein structures—to assess structural validity at scale, we mapped crosslinks onto computationally predicted structures, where intra-protein crosslinks were mapped to top-scoring AlphaFold2^32^ monomeric structures (**Fig. 1H**), and inter-protein crosslinks were mapped to AlphaFold-Multimer^48^ complex predictions. These computationally mapped crosslinks were utilized downstream for AlphaLink structural refinement (**Fig. 1I**).

### Distance constraint violations reveal potential tumor-specific conformations

Using these AlphaCross-XL-derived Cα-Cα distances, we assessed structural constraint satisfaction and identified potential conformational changes. Crosslinks were mapped onto top-scoring AlphaFold2 monomer or AlphaFold-Multimer structures, as appropriate, with C_α_-C_α_ distances computed between each crosslinked residue pair. DSSO crosslinks exhibited narrower distance distributions (median: 15.06 Å, n=1,016) compared to PhoX crosslinks (median: 17.38 Å, n=2,662), likely reflecting differences in spacer arm chemistry, though both showed similar overall distribution profiles (**Fig. 2A-B**).

**Figure 2.**
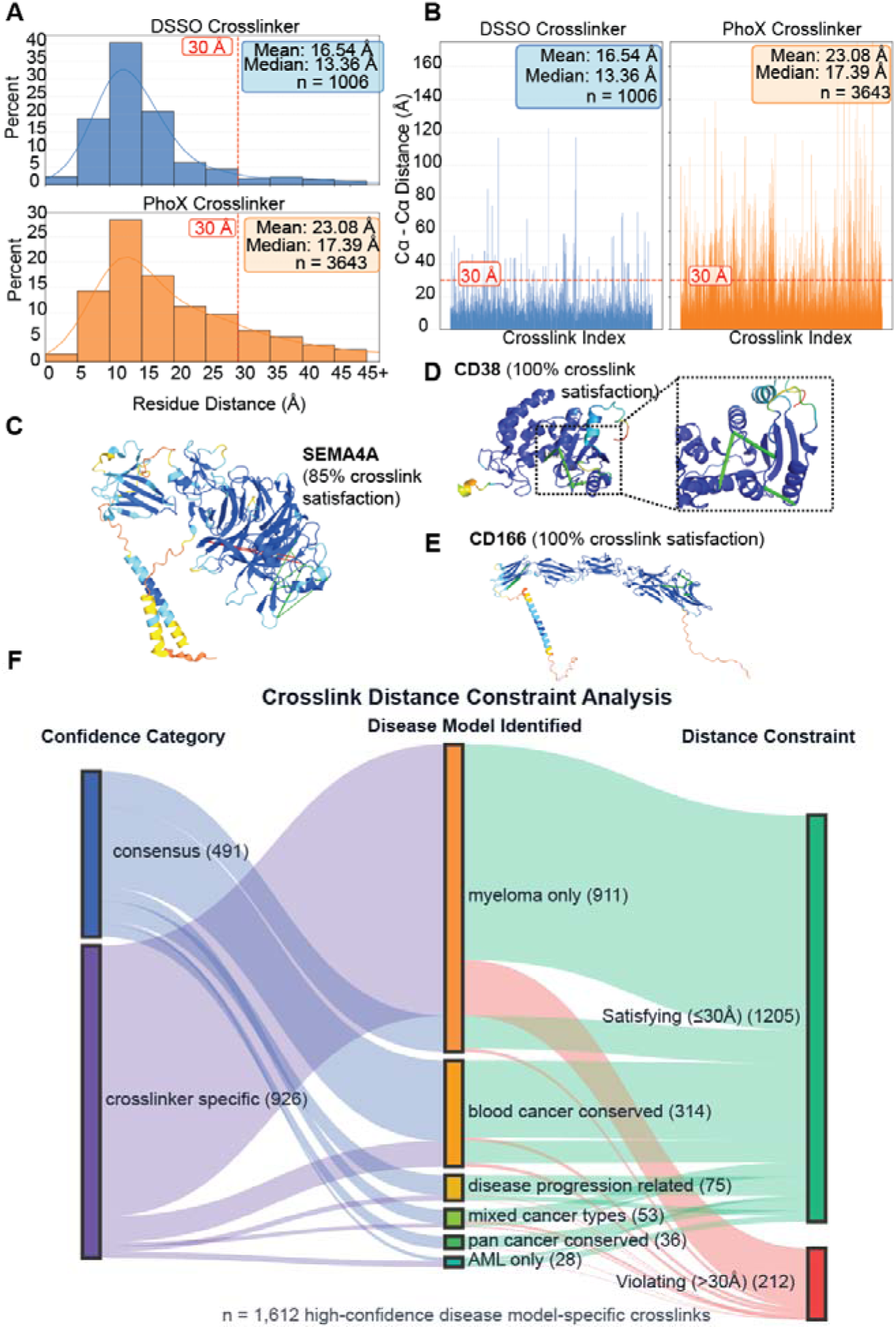
Structural Validation and Identification of Disease-Specific Conformations. **(A)** Histograms of residue distances for crosslinks detected by DSSO (top, blue) and PhoX (bottom, orange) crosslinkers. Histogram bins are 5 Å intervals with the final bin capturing all distances ≥45 Å. Kernel density estimation curves (solid lines) show smoothed distributions. The red dashed line at 30 Å indicates the maximum expected Cα-Cα distance for valid crosslinks, based on the combined reach of the crosslinker and lysine side chains. DSSO crosslinks show tighter distance distributions (Mean: 16.52 Å, Median: 15.06 Å, n = 1016) compared to PhoX (Mean: 22.08 Å, Median: 17.38 Å, n = 2662), with most crosslinks falling below the 30 Å threshold, validating structural accuracy. **(B)** Individual residue distances for DSSO (left, blue) and PhoX (right, orange) crosslinks plotted by crosslink index, where each bar represents a single crosslink observation. The red dashed line at 30 Å marks the theoretical distance constraint. Both crosslinkers show predominantly valid distances below 30 Å, with PhoX displaying more variability and a greater proportion of longer-distance crosslinks. **(C-E)** Example proteins with both satisfied and violated crosslinks mapped to structure, illustrating conformational difference concept. **(F)** Classification and structural validation of high-confidence disease-specific crosslinks (n = 1,612). Flow diagram shows crosslink distribution from validation category (left: consensus vs. crosslinker specific) through disease classification (middle) to distance constraint outcome (right: satisfying ≤30Å vs. violating >30Å based on AlphaFold models). Flow width indicates crosslink count. Consensus crosslinks show 5.9% violation rate compared to 16.9% for crosslinker-specific links. Myeloma-specific crosslinks (n = 911) represent the largest disease category, with 176 distance violations potentially indicating malignancy-associated conformational changes.

The majority of crosslinks from both chemistries satisfied the theoretical 30 Å distance constraint, validating structural accuracy. However, a subset of crosslinks exceeded the 30 Å threshold, indicating potential conformational differences between experimental and predicted states. These distance constraint violations could indicate conformational differences between experimentally captured states and computationally predicted reference structures. Example proteins showed both satisfied and violated crosslinks (**Fig. 2C-E**), suggesting possible regional conformational remodeling.

For proteins with multiple experimental structures, we applied xlms-tools^49^ to score structures against our crosslink data (**Fig. S4)**. One such instance displayed within these datasets is the sodium/potassium-transporting ATPase, which is known to take two distinct conformations based on which substrate is bound: E1·3Na and E1·3Na·ATP (PBD: 7E1Z) or E2·[2K] (PBD: 7E20). Within the THP-1 samples crosslinked with DSSO, seven crosslinks were identified on the Na+/K+ ATPase and subsequently mapped onto the three separate cryo-EM derived structures. When the crosslinks are mapped onto the E1·3Na and E1·3Na·ATP structures, there is one distance violation identified on both conformations, with the C_α_-C_α_ distances being 34.38 Å and 34.55 Å respectively. However, this crosslink distance decreases to 20.20 Å when mapped onto the E2·[2K] conformation, therefore being well-within the distance limit. This analysis implies the Na^+^/K^+^ ATPase is primarily in its 2K-bound conformation on the THP-1 cell line and demonstrates that structural surfaceomics may be able to distinguish previously-characterized membrane protein conformations.

Returning to our broader dataset, we classified 1,612 “disease model-specific” crosslinks (i.e. crosslinks only appearing in a dataset for one indication represented by our tumor models, whether myeloma, AML, or prostate cancer) by validation category—consensus (both DSSO and PhoX) versus crosslinker-specific (one chemistry)—and assessed distance constraint satisfaction (**Fig. 2F**). Consensus crosslinks showed a 5.9% violation rate compared to 16.9% for crosslinker-specific crosslinks, indicating higher confidence for consensus observations. Multiple myeloma yielded the most disease model-specific crosslinks (n=911), with 176 violations potentially indicating conformational changes. Across all tumor models explored here, we identified 212 high-confidence violations representing candidate conformation-specific epitopes.

### Structural characterization of conformational epitopes

To identify structural features associated with conformational remodeling, we performed domain enrichment analysis on the 212 distance-violating crosslinks. We focused on proteins with well-structured regions (AlphaFold prediction score pLDDT >70) to avoid analysis of highly dynamic and/or intrinsically disordered regions (**Fig. 3A-B**). Cancer-specific crosslinks were significantly enriched in regions with loop/coil secondary structure (Odds Ratio, OR=1.21, FDR<0.05) and alpha helices (OR=1.67 for AML) (**Fig. 3C**). More broadly, immunoglobulin-like domains (59 proteins), repeat domains including ARM-type and leucine-rich repeats (301 proteins), and signaling/interaction domains such as PH-like and SH3 domains (383 total proteins) were enriched in the datasets (**Fig. 3D**), suggesting that these structural elements are particularly susceptible to conformational changes in malignancy and/or particularly likely to drive errors in the AlphaFold prediction. Many of these proteins consisted of less than 10 domains, which argues against multi-domain complexity as the primary driver of the observed violations (**Fig. 3E**). This distribution most affected proteins that have architecturally simple folds for which AlphaFold predictions are typically most reliable. The small subset of highly complex proteins at the tail of the distribution (detailed in **Fig. 3F**, including LRIG family members, obscurin, and the Notch receptor family with 29–40 domains each) represents cases where prediction uncertainty may be a contributing factor and warrant additional experimental scrutiny. Functional analysis of proteins revealed enrichment in regulation of actin cytoskeleton, cell adhesion molecules, and endocytosis pathways (**Fig. 3F**), implicating cell adhesion, signal transduction, and chromatin organization in cancer-associated conformational remodeling.

**Figure 3.**
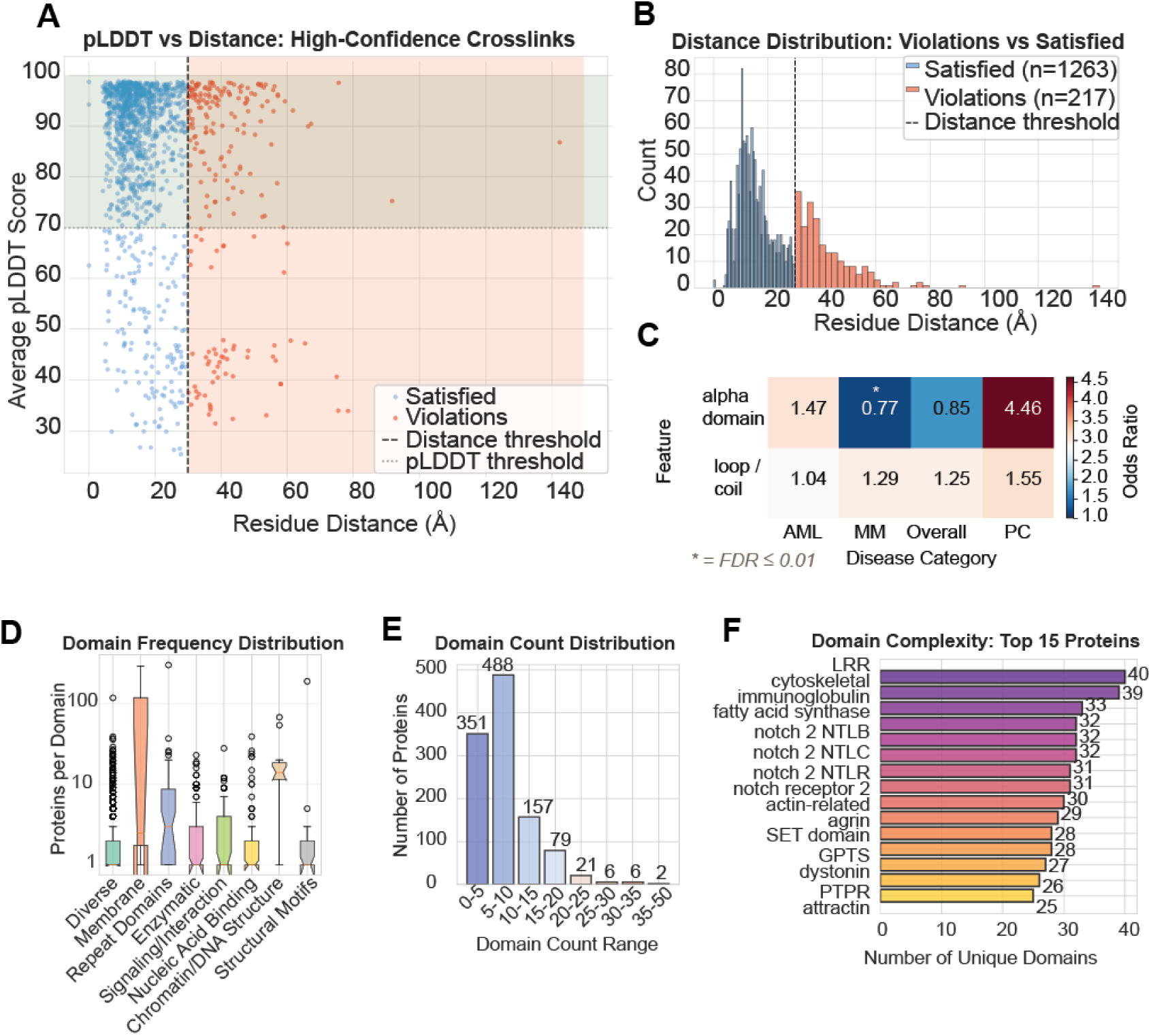
Domain and Structural Enrichment Analysis of Cancer-Specific Crosslinks. **(A)** pLDDT versus Cα–Cα distance for all crosslinks; satisfied crosslinks (blue) cluster in the high-confidence, short-distance quadrant while violations (red) span confidence levels. The 35 Å distance and pLDDT = 70 thresholds define regions for downstream filtering. **(B)** Distance distributions of satisfied and violated crosslinks (extending >100 Å) show a bimodal distribution pattern. **(C)** Heatmap of secondary structure enrichment in distance-violating crosslinks; alpha-helical regions are enriched in AML (OR = 1.67) and loop/coil regions broadly across disease types (OR = 1.21–1.54, FDR < 0.05). **(D)** Box plots of proteins per domain type by functional category (log scale); repeat and signaling/interaction domains show the broadest distributions. **(E)** Domain count distribution among violating proteins peaks at 0–5 unique domains (n ≈ 430), demonstrating that most violating proteins are structurally compact—arguing against multi-domain complexity as the primary driver of violations. **(F)** Top 15 proteins by unique domain count (range: 25–40), including LRIG family (40 domains), obscurin (39), and Notch receptor 2 (31–32); these high-complexity cases warrant additional validation before violations are interpreted as conformational evidence.

To distinguish genuine conformational changes from prediction uncertainty, we analyzed predicted Local Distance Difference Test (pLDDT) scores and Predicted Aligned Error (PAE) metrics at crosslinked residues. Distance-violating crosslinks occurred predominantly in high-confidence regions (median pLDDT: 89.9, n=217) with variable PAE values (**Fig. S6**), indicating that violations may reflect genuine conformational changes rather than prediction uncertainty. In contrast, satisfied crosslinks showed similar confidence metrics (median pLDDT: 91.9, p=0.006), confirming that AlphaFold structures provide reliable reference models for identifying cancer-specific conformations. 27.2% of violating crosslinks mapped to flexible or disordered regions (pLDDT <70), suggesting these violations may reflect AlphaFold prediction uncertainty rather than cancer-specific conformations (**Fig. 4A-D**). Filtering for high-confidence violations (pLDDT >70) yielded 158 prioritized crosslinks for structural refinement, including CD48 in multiple myeloma and integrin α1/β4 in AML, which we investigate in more detail below.

**Figure 4.**
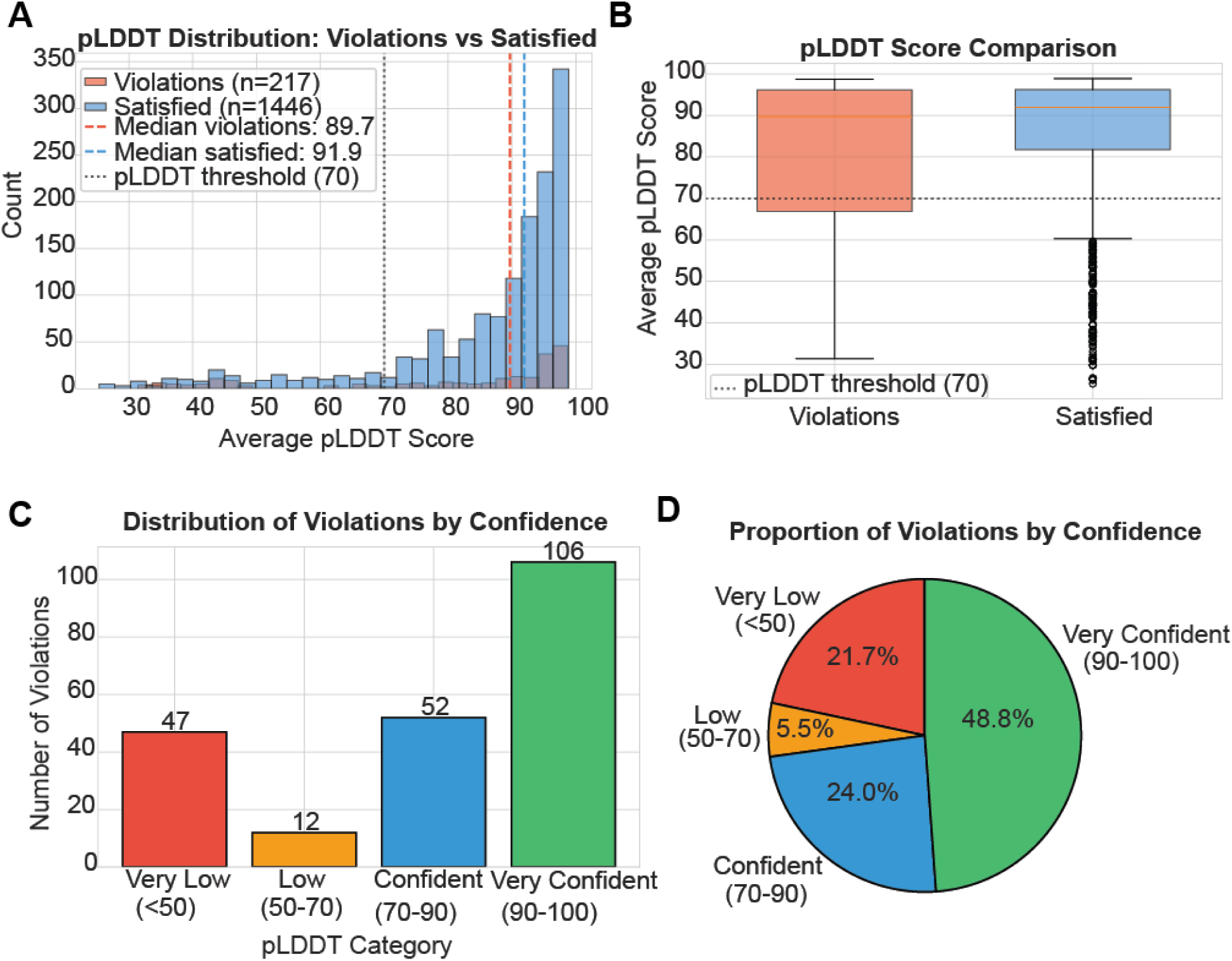
AlphaFold confidence analysis validates structural accuracy of disease-specific crosslinks. **(A)** Distribution of average pLDDT scores for crosslinked residue pairs in distance-violating (red, n=217) versus satisfied (blue, n=1,446) crosslinks. Vertical dashed lines indicate median pLDDT scores (violations: 89.7; satisfied: 91.9). Gray dotted line marks the pLDDT threshold of 70, below which regions are considered flexible or disordered. Both populations show predominantly high confidence scores (>70), with distance violations occurring primarily in well-predicted regions rather than disordered loops, supporting genuine conformational differences over prediction artifacts. **(B)** Box plot comparison of pLDDT scores between violating and satisfied crosslinks. Both groups exhibit high median confidence (violations: 89.7; satisfied: 91.9; Mann-Whitney U test, p=0.006), confirming AlphaFold structures as reliable reference models. Gray dotted line indicates pLDDT threshold of 70. Only 27.2% of violations (59/217) mapped to flexible regions (pLDDT <70), with 72.8% (158/217) representing high-confidence violations that likely reflect cancer-specific conformational changes rather than prediction uncertainty. Individual outliers are shown as black points. **(C)** Bar chart showing the number of distance-violating crosslinks stratified by AlphaFold pLDDT confidence category. Violations are observed across all confidence bins, including 106 violations in the “Very Confident” range (pLDDT 90–100), 52 in “Confident” (70–90), 12 in “Low” (50–70), and 47 in “Very Low” (<50). The enrichment of violations in high-confidence regions further indicates that these discrepancies are not confined to poorly predicted or disordered segments, but instead often arise in well-modeled structural contexts. **(D)** Pie chart summarizing the proportion of distance-violating crosslinks within each AlphaFold pLDDT confidence category. Nearly half of all violations (48.8%) occur in “Very Confident” regions (90–100), with an additional 24.0% in “Confident” (70–90), 5.5% in “Low” (50–70), and 21.7% in “Very Low” (<50). The substantial fraction of violations in high-confidence regions supports the interpretation that many represent bona fide disease-specific conformational changes rather than artifacts of uncertain structure prediction.

### Case study 1: Multiple myeloma-specific CD48 conformation

To investigate whether distance violations represent bona fide conformational changes, we performed detailed structural modeling of CD48, a cell surface receptor with multiple myeloma-specific crosslinks. CD48 is one of the most highly expressed myeloma surface antigens and an attractive therapeutic target, but “on target, off tumor” hematologic toxicity of a CD48-targeting antibody-drug conjugate led to clinical trial discontinuation due to CD48 expression on normal hematopoietic cells^50,51^. As one of the most highly expressed myeloma surface antigens^40^, CD48 appears to be a highly promising therapeutic target if a tumor-specific epitope or conformation can be identified for targeting the tumor while sparing normal cells.

Analysis of 10 crosslinks detected consistently across both myeloma cell lines (AMO-1 and MM.1S) revealed a mixed pattern: four crosslinks exceeded the 30 Å threshold when mapped to AlphaFold structure (Residue 80-114: 30.6 Å; 98-114: 34.6 Å; 124-137: 36.3 Å; 75-133: 42.6 Å), while six satisfied distance constraints (**Table 1**, **Fig. 5A**). All crosslinked residues showed high AlphaFold confidence (pLDDT >90), indicating violations reflect genuine conformational changes rather than prediction uncertainty. Violations clustered at the receptor binding loop (residues 114-117) and interdomain linker (residues 128-135) (**Fig. 5B**).

**Figure 5.**
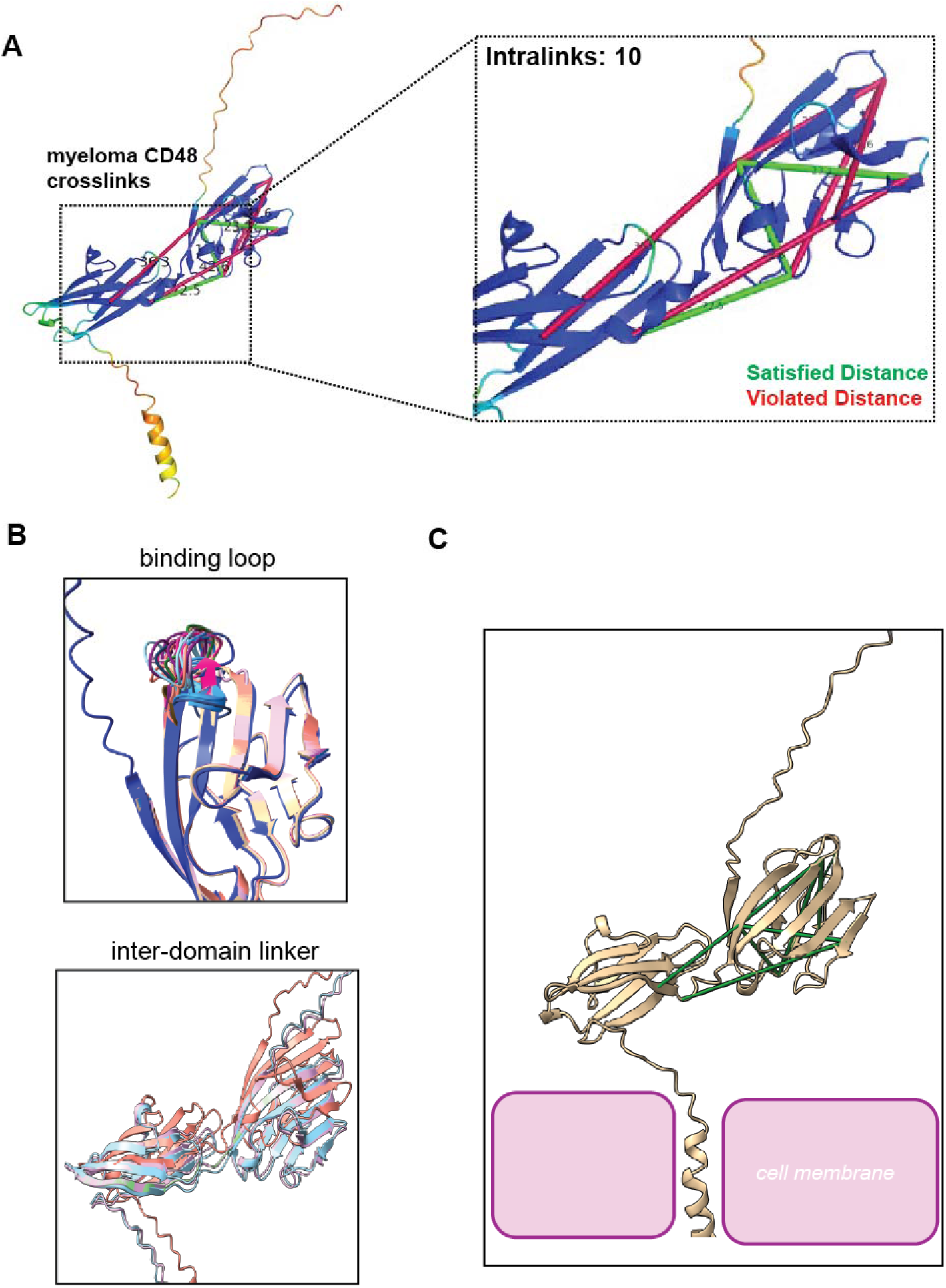
Crosslink-guided Structural Refinement Reveals a Multiple Myeloma-specific CD48 Conformation. **(A)** CD48 crosslinks detected across myeloma (AMO-1, MM.1S) and AML (NOMO-1) cell lines mapped onto the AlphaFold2-predicted structure, color-coded by distance constraint status: satisfied (≤30 Å, blue) or violated (>30 Å, red). Four myeloma-specific crosslinks exceed the 30 Å threshold. All crosslinked residues show high AlphaFold confidence (pLDDT >90). **(B)** Structural localization of crosslink violations on CD48. Violated crosslinks cluster at the receptor binding loop (residues 114-117) and interdomain linker (residues 128-135), both in high-confidence structural regions (pLDDT >90), indicating these violations reflect genuine conformational differences rather than prediction uncertainty. **(C)** Crosslink-informed MODELLER refinement of CD48. The refined model (green) adopts a compact “closed” conformation satisfying all 10 detected crosslinks, with repositioning of the receptor binding loop and reorientation of the interdomain linker relative to the canonical AlphaFold reference structure (gray). This myeloma-specific conformation satisfies the distance constraints of the crosslinker.

**Table 1:** XL-MS data for CD48 is detected in myeloma (AMO1, MM1.S) and AML (NOMO1).

| CD48 Crosslink<br>(Residue Pair) | Spectral Count | Detected In (Datasets) | Cα-Cα<br>Distance (Å) | pLDDT<br>(Res. 1) | pLDDT<br>(Res. 2) |
| --- | --- | --- | --- | --- | --- |
| Lys75-Lys114 | 8 | Myeloma (PhoX, n=2) | 19.25 Å | 92.58 | 96.75 |
| Lys80-Lys114 | 4 | Myeloma (PhoX, n=2) | 30.65 Å | 92.58 | 93.5 |
| Lys114-Lys124 | 14 | AML (PhoX, n=1); Myeloma<br>(PhoX, n=2) | 27.54 Å | 96.66 | 92.58 |
| Lys124-Lys137 | 3 | Myeloma (PhoX, n=2) | 36.35 Å | 96.66 | 95.87 |
| Lys75-Lys124 | 6 | Myeloma (PhoX, n=2) | 23.07 Å | 96.66 | 96.75 |
| Lys75-Lys133 | 5 | Myeloma (PhoX, n=2) | 42.59 Å | 95.57 | 96.75 |
| Lys98-Lys114 | 12 | Myeloma (PhoX, n=2) | 34.63 Å | 97.72 | 92.58 |
| Lys98-Lys124 | 6 | Myeloma (PhoX, n=2) | 17.02 Å | 97.72 | 96.66 |
| Lys98-Lys133 | 3 | Myeloma (PhoX, n=2) | 22.51 Å | 97.72 | 95.57 |
| Lys75-Lys98 | 6 | Myeloma (PhoX, n=2) | 20.80 Å | 97.72 | 96.75 |

We refined the CD48 structure using MODELLER^27,52,53^, a comparative protein structure modeling program that generates conformations by satisfying user-defined spatial restraints. We utilized our experimental XL-MS data with experimental crosslink restraints, keeping residues with satisfied crosslinks static and allowing flexibility in violation regions^26,28^ (**Fig. 5C**). The crosslink-informed model repositioned the binding loop and reoriented the interdomain linker, producing a compact “closed” conformation that satisfied all 10 crosslinks. If a conformation-selective binder could be developed, this myeloma-specific conformation could enable antibodies to selectively target tumor CD48 while sparing normal cells. Such a therapeutic would potentially reduce on-target/off-tumor toxicity and establish CD48 as a priority candidate for conformation-selective immunotherapy.

### AML-specific integrin heterodimer conformations

To further validate the AML-specific LFA-1 heterodimer conformation from our previous work with increased depth of coverage, we analyzed the integrin conformational landscape detected across AML cell line models (**Fig. 6A**, **Table 2**). In previous work, the open conformation of LFA-1 was identified on DSSO-labeled NOMO1 structural surfaceome^18^. Here, we recapitulate those findings across additional AML models — THP-1 and MV411 — revealing expanded crosslink coverage with both DSSO and PhoX labeling. Across AML cell lines, we detected 13 inter-protein crosslinks connecting αL and β2 subunits, with all 13 specific to AML models and absent from myeloma, prostate cancer, and normal PBMC datasets (**Fig. 6B**, **Table 2**).

**Figure 6.**
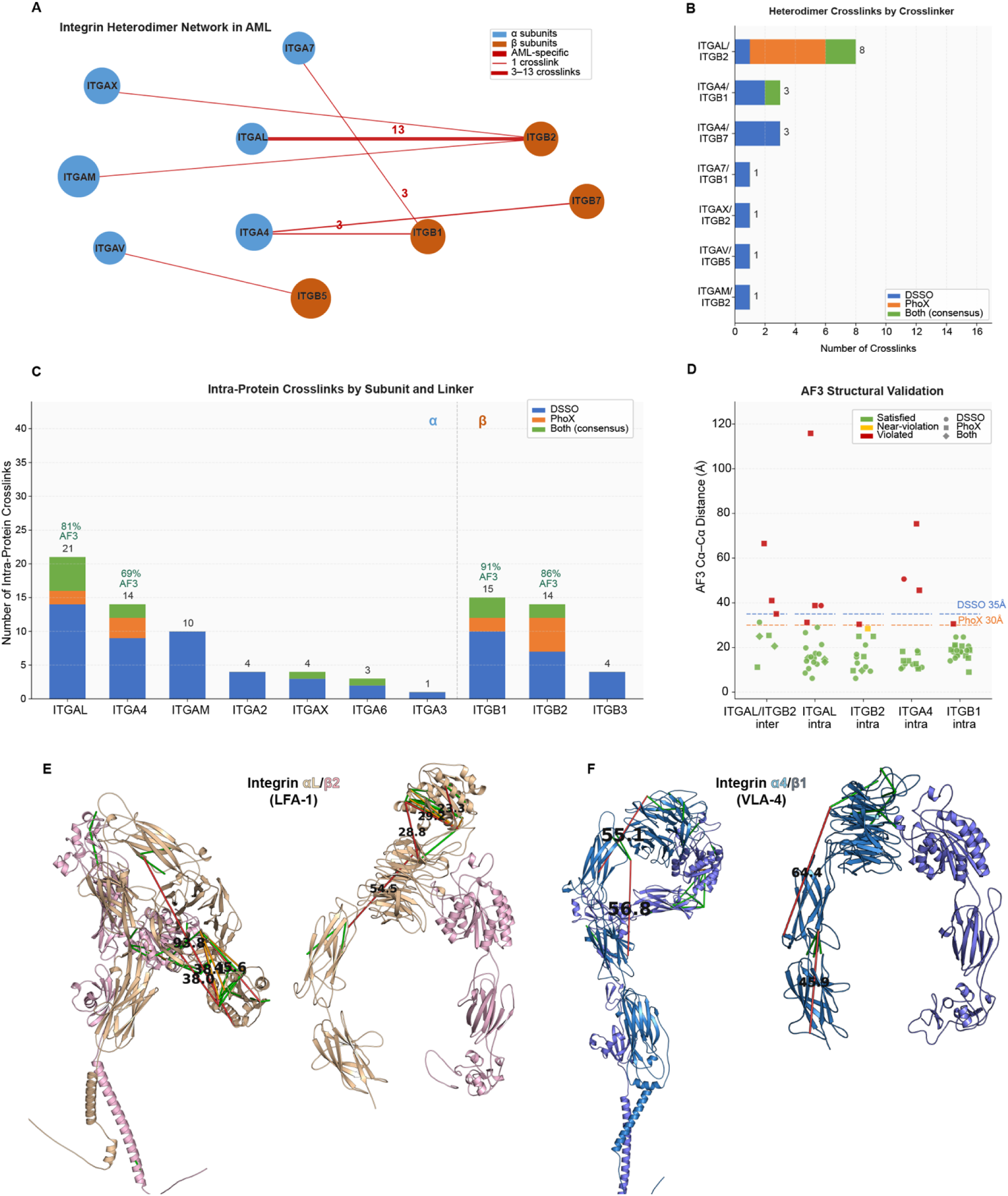
AML-specific Integrin Heterodimer Crosslinks and Conformational Validation. **(A)** Network diagram illustrating the landscape of integrin α and β subunit inter-protein crosslinks detected across AML cell line models. Node size reflects the number of crosslinked subunit partners; edge thickness indicates crosslink count. All inter-subunit crosslinks shown are AML-specific. **(B)** Inter-protein crosslink counts by integrin heterodimer pair and crosslinker (DSSO, PhoX, or both). Consensus crosslinks independently detected with both crosslinkers are highlighted in green. The αL/β2 (LFA-1) pair harbors the largest inter-subunit crosslink coverage (n = 13). **(C)** Intra-protein crosslink counts by integrin subunit and crosslinker, showing the distribution of detected DSSO, PhoX, and consensus crosslinks across individual α and β subunits. **(D)** Cα–Cα distance plot for all crosslinks evaluated against AF3 heterodimer structures of LFA-1 (ITGαL/ITGβ2) and VLA-4 (ITGα4/ITG β1). Each point represents a unique lysine pair; shape indicates crosslinker (DSSO, 35 Å threshold; PhoX, 30 Å threshold); color indicates satisfaction status (green: satisfied; yellow: near-limit; red: violated). Dashed lines denote crosslinker thresholds. For LFA-1, 5/8 inter-subunit pairs and 30/35 intra-subunit pairs were satisfied; for VLA-4, 34/38 intra-subunit pairs were satisfied. **(E)** AF3 model of LFA-1 (Integrin αL/β2) with inter- and intra-subunit crosslinks visualized. Crosslinks are colored by distance status: green (satisfied), yellow (near-limit), red (violated). The three violated inter-subunit crosslinks, including K293(αL)–K148(β2) at 66.5 Å, localize to the αL–β2 headpiece interface. **(F)** AF3 model of VLA-4 (Integrin α4/β1, closed conformation) with intra-subunit crosslinks visualized. Violated crosslinks, including K526(α4)–K178(α4) at 75.4 Å, span the hybrid domain and β-propeller head — contacts incompatible with the closed-state model. **(G)** Open extended conformation of VLA-4 generated by homology modeling, with crosslinks overlaid. Transition to the open conformation reduces distance violations for crosslinks spanning the α4 headpiece, supporting a conformational state distinct from the AF3 closed-state structure detected on the AML cell surface.

**Table 2:** Crosslink data for Integrins and Integrin complexes.

| Integrin Pair | Total XL | Type | AML-specific | High Conf. | Mean Dist. AF2 (Å) | AF3 Mean Dist. (Å) | AF3 Violations | xlms-tools score DSSO | xlms-tools score PhoX |
| --- | --- | --- | --- | --- | --- | --- | --- | --- | --- |
| ITGAL (intra) | 33 | Intra | 32 | 6 | 23.5 | 23.6 | 4 | 0.617 | 0.070 |
| ITGB1 (intra) | 31 | Intra | 21 | 6 | 18.2 | 18.8 | 1 | 0.792 | 0.269 |
| ITGB2 (intra) | 26 | Intra | 23 | 1 | 16.6 | 17.0 | 1 | 0.617 | 0.070 |
| ITGA4 (intra) | 18 | Intra | 10 | 3 | 19.4 | 21.6 | 3 | 0.792 | 0.269 |
| ITGAM (intra) | 15 | Intra | 13 | 1 | 26.4 | — | — | — | — |
| ITGAL/ITGB2 | 8 | Inter | 8 | 0 | — | 35.1 | 3 | 0.617 | 0.070 |
| ITGA4/ITGB7 | 3 | Inter | 3 | 0 | — | — | — | — | — |
| ITGA4/ITGB1 | 3 | Inter | 3 | 0 | — | 21.6 | — | — | — |

Beyond the extensively crosslinked LFA-1 complex, our surfaceomics approach identified crosslinks for six additional integrin heterodimers, revealing a broader landscape of integrin conformational states in AML (**Fig. 6B, C**). We detected three inter-protein crosslinks each for VLA-4 (CD29d, ITG α4/β1) and LPAM-1 (CD49d, ITG α4/β7), both of which mediate AML cell adhesion to bone marrow stroma and endothelium. All crosslinks for these additional heterodimers were AML model-specific and absent from myeloma, prostate cancer, and normal hematopoietic cells. We then used AlphaFold3 heterodimer structural predictions of Integrin αL/β2 and α4/β1 to compute structural distances for the detected crosslinks (**Fig. 6D**). For LFA-1 (αL/β2), we evaluated 8 unique inter-subunit lysine pairs against the AF3 closed-conformation model: 5/8 were satisfied (63%) and 3 violated, with the largest discrepancy at K293(αL)–K148(β2), spanning 66.5 Å against the 30 Å PhoX threshold. Intra-subunit crosslinks showed strong overall agreement for both αL (17/21 satisfied, 81%; mean Cα–Cα distance 23.6 Å) and β2 (13/14 satisfied, 93%; mean 17.0 Å). Structural validation with xlms-tools yielded a DSSO average crosslink score of 0.617 for the LFA-1 AF3 model, consistent with the inter-subunit violations concentrated at the headpiece interface (**Fig. 6E**). These violations are consistent with our previous observations and reflect crosslinks captured in the extended, open-headpiece conformation that is incompatible with the closed-state AF3 structure.

For VLA-4 (α4/β1), intra-subunit crosslinks showed strong compatibility with the AF3 closed-conformation model for both α4 (13/16 satisfied, 81%; mean 21.6 Å) and β1 (21/22 satisfied, 95%; mean 18.8 Å), with an overall xlms-tools DSSO score of 0.792 (**Fig. 6F**). However, three α4 intra-crosslinks were violated in the closed-state model, including a particularly large discrepancy at K526(α4)–K178(α4) (75.4 Å, PhoX threshold 30 Å) — a contact spanning the hybrid domain and the β-propeller head that is incompatible with the compact, closed-headpiece AF3 structure. To assess whether the open conformation of VLA-4 better accommodates these crosslinks, we generated a homology model of the extended α4/β1 headpiece. This open-state model reduced the distance violation for the inter-domain crosslinks spanning the headpiece (**Fig. 6G**), consistent with a conformational deviation from the AF3 closed structure captured by crosslinking. Flow cytometry using the conformation-sensitive CD29d antibody, which selectively recognizes the extended, active β1 integrin headpiece, confirmed expression of the active VLA-4 conformation on AML cell lines, with strongest signal on THP-1 cells (**Fig. S8**). Together, these crosslinking, structural modeling, and flow cytometry data support the presence of distinct, open-headpiece integrin quaternary conformations in AML, consistent with conformational remodeling of both LFA-1 and VLA-4 on the leukemic cell surface.

### XL-MS integrative modeling across the surfaceome

Beyond identifying distance violations, we sought to leverage our experimental crosslink data to directly improve protein structure prediction through XL-MS integrative modeling. In this paradigm, experimentally derived distance restraints are incorporated as soft constraints during the prediction process, guiding the model toward conformations consistent with the observed crosslinking data. We assessed this strategy by supplying crosslink-derived distance restraints to AlphaLink^33^, an AlphaFold-derived network trained to incorporate XL-MS contacts as structural priors. Across 278 surfaceome proteins encompassing 2,871 crosslinks, AlphaLink produced a statistically significant reduction in Cα–Cα crosslink distances relative to AlphaFold2 (mean 23.2 → 22.9 Å; Wilcoxon signed-rank p = 3.3 × 10 □¹ □), though the global effect size was modest (Wilcoxon signed-rank test rank-biserial, r = −0.15; **Fig. 7A, B**).

**Figure 7.**
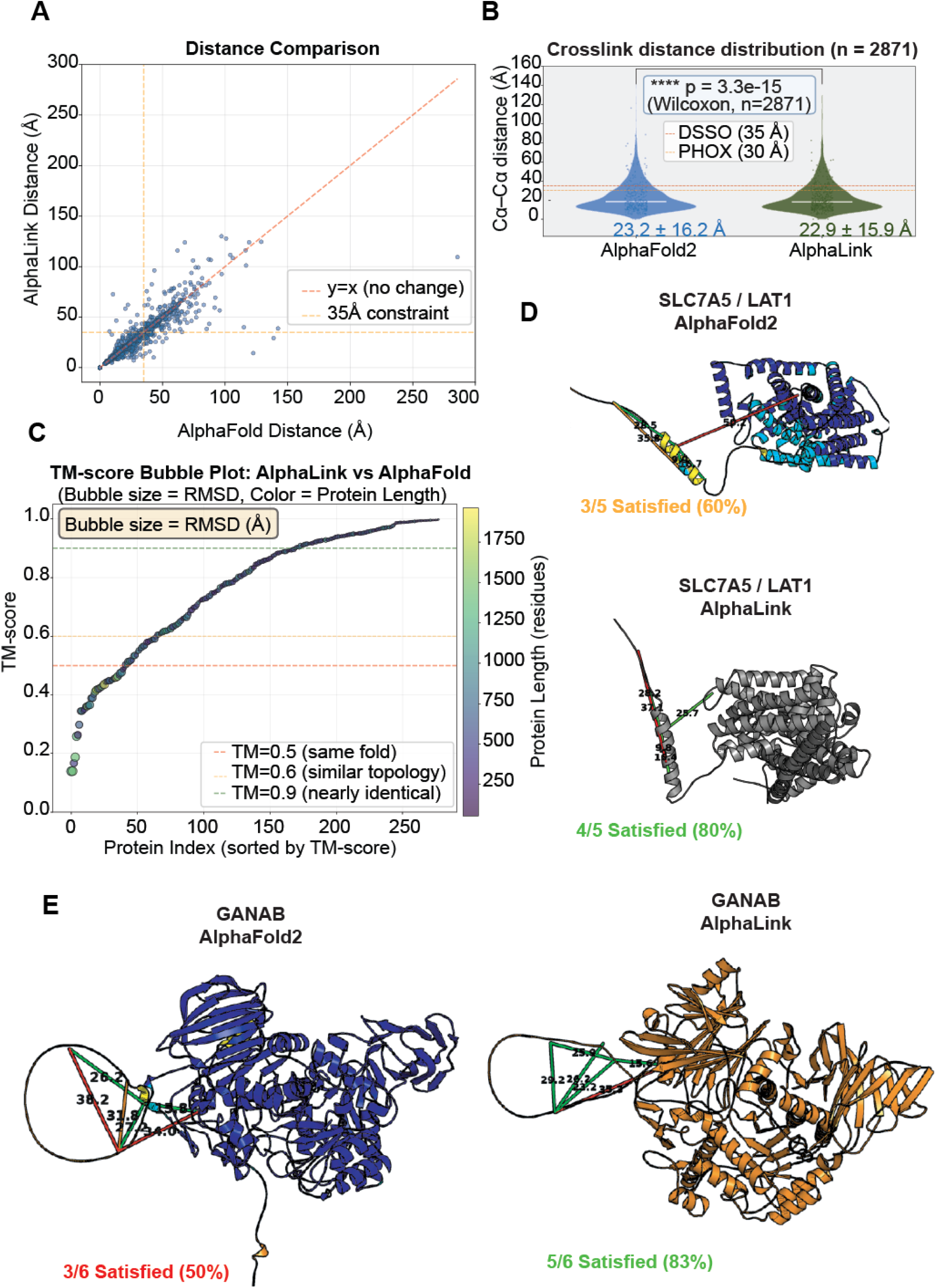
AlphaLink Integrative Modeling Performance Across the Surfaceome. **(A)** Pairwise scatter plot of Cα–Cα crosslink distances computed from AlphaLink vs. AlphaFold2 models (n = 2,871 crosslinks across 278 proteins). The y = x diagonal (pink dashed) indicates no change; the orange dashed lines denote the 35 Å DSSO constraint. The majority of crosslinks cluster tightly around the diagonal, indicating that AlphaLink preserves the global fold while making targeted local adjustments. **(B)** Violin plots comparing the crosslink Cα–Cα distance distributions between AlphaFold2 (mean = 23.2 ± 36.2 Å) and AlphaLink (mean = 22.9 ± 35.9 Å) across all 2,871 crosslinks. Horizontal dashed lines indicate the DSSO (35 Å) and PhoX (30 Å) distance thresholds. AlphaLink produces a modest but statistically significant reduction in crosslink distances (Wilcoxon signed-rank test, W = 2,375,185, p = 3.3 × 10 ¹ ; effect size r = −0.15). **(C)** TM-score bubble plot for 278 surfaceome proteins with AlphaLink predictions, sorted by ascending TM-score (AlphaLink vs. AlphaFold2). Bubble size reflects Cα RMSD; bubble color reflects protein chain length. Reference lines are shown at TM = 0.5 (same fold threshold, red), TM = 0.6 (similar topology, green dashed), and TM = 0.9 (near-identical, green dashed). The majority of proteins achieve TM > 0.9, indicating that AlphaLink largely preserves the AlphaFold2 fold; a low-TM tail (left) identifies proteins where crosslink restraints drive substantive conformational remodeling, including candidate targets LFA-1 and CD48. **(D)** Representative structural comparisons for two proteins showing clear AlphaLink improvement. SLC7A5/LAT1 (top): AlphaFold2 satisfies 3/5 crosslinks (60%); AlphaLink satisfies 4/5 (80%). GANAB (bottom): AlphaFold2 satisfies 3/6 crosslinks (50%); AlphaLink satisfies 5/6 (83%). Backbone ribbon is colored by pLDDT confidence score; crosslinks are shown as dashed lines colored by satisfaction status (green: satisfied, red: violated).

At the binary level, AlphaLink rescued 57 crosslinks from violated to satisfied while worsening 35 other crosslinks (McNemar χ² = 4.79, p = 0.029), confirming a net improvement in structural crosslink compatibility. Notably, crosslink improvement was independent of the magnitude of structural change: TM-score between AlphaLink and AlphaFold2 models showed no correlation with per-protein crosslink distance improvement (Spearman r = −0.003), indicating that AlphaLink refines local inter-residue contacts at specific sites rather than driving global conformational remodeling. Exemplary improvements were observed for SLC7A5/LAT1 (AlphaFold2: 3/5 satisfied, 60% → AlphaLink: 4/5, 80%) and GANAB (3/6, 50% → 5/6, 83%; **Fig. 7D**), proteins for which the crosslink-derived restraints guided the model toward more compact, experimentally consistent local geometries.

In contrast, for our two candidate cell-surface targets — LFA-1 (αL/β2) in AML models and CD48 in myeloma models — AlphaLink models exhibited globally perturbed topologies, with markedly reduced TM-scores relative to the original AlphaFold2 structures despite consistent and reproducible underlying crosslink data (**Fig. 7C**). These proteins fall within the low-TM tail of the distribution, where crosslink restraints appear to conflict with the AlphaFold prediction, driving the model away from the canonical topology. We interpret this failure as a consequence of the membrane environment and cancer specific cell-surface dynamics that constrain these proteins *in vivo* but are absent from AlphaLink’s training paradigm. This failure mode is itself informative: it delineates the current boundaries of XL-MS integrative modeling tools and motivates the development of surface-context-aware structural prediction architectures. For these targets, the crosslinking data are better interpreted through explicit conformational modeling against known structural states, as demonstrated by the homology modeling and flow cytometry validation for VLA-4 (**Fig. 6F, G; Fig. S7**), rather than through unconstrained deep learning refinement.

## DISCUSSION

This study establishes structural surfaceomics as a systematic approach to identify cancer-specific protein conformations, moving beyond traditional expression-based target discovery^12,13^. By integrating crosslinking mass spectrometry with surface biotinylation across multiple cancer models, we identified 212 potentially tumor-specific protein structure alterations based on distance constraint violations relative to AlphaFold predictions in well-folded, high confidence regions. These findings support the notion that malignancy-driven conformational changes may be not rare exceptions but rather systematically detectable features of the cancer surfaceome, with 13% of disease model-specific crosslinks violating predicted structural constraints. The identification of cancer-specific violating crosslinks in well-characterized proteins like CD48 and integrin α4/β1suggests that conformational epitopes represent an underexplored therapeutic space, particularly relevant as many traditional, expression-based surface targets reach clinical saturation.

The conformational changes detected through structural surfaceomics likely reflect fundamental alterations in cancer cell biology. The AML-specific stabilization of integrin α1/β4 in its active conformation exemplifies how cancer cells remodel surface protein complexes to support malignant phenotypes. Constitutive stabilization of integrin α1/β4 in the active conformation has important implications for AML pathogenesis. Active integrins mediate high-affinity binding to extracellular matrix ligands, promoting AML cell adhesion to the bone marrow microenvironment—a key mechanism of chemotherapy resistance through cell adhesion-mediated drug resistance (CAM-DR). In fact, active integrin α4/β1 represents a previously explored target in AML and solid tumors using a small molecule inhibitor in several reported clinical trials but has not progressed past Phase II^54,55^. Similarly, the multiple myeloma-specific CD48 conformation, characterized by interdomain flexibility and binding loop repositioning, potentially reflects altered cellular signaling, microenvironment interactions, or mechanical stresses unique to the malignant state. These conformational shifts may arise from aberrant post-translational modifications, altered protein-protein interactions, or changes in membrane organization driven by oncogenic signaling. Future studies will be required to investigate these possibilities.

The breadth of AML-specific integrin crosslinks is also mechanistically notable. Detection of multiple β2-containing complexes (LFA-1, Mac-1, and αXβ2) is consistent with the leukocyte-restricted expression of β2 integrins and their established roles in immune trafficking and AML blast migration. Similarly, detection of both α4-containing heterodimers (VLA-4 and LPAM-1) resonates with the well-established role of α4 integrins in AML cell adhesion to the bone marrow microenvironment. These findings raise the possibility that AML may involve coordinated conformational or expression changes across functionally related integrin families, potentially reflecting the altered adhesion that characterizes leukemic progression and drug resistance. Future studies correlating per-cell line crosslink patterns with quantitative protein abundance measurements will be needed to disentangle conformational from expression-level contributions to these observations.

Beyond the specific examples presented here, our results suggest that conformational epitopes represent a fundamentally distinct class of immunotherapy target, one not previously accessible through expression-based proteomics approaches. These also include surfaceome mapping that only provides catalogues of surface bound proteins and dynamic surfaceome maps without structural information^56,57^. Traditional surface targets are typically defined by differential abundance — tumor enrichment versus normal tissue expression — but this gradient rarely achieves the clean binary separation needed for a wide therapeutic window. Conformational epitopes, by contrast, operate on a different logic: a cancer-associated protein shape is either present or absent on a given cell, independent of expression level. This binary character is clinically meaningful, because it permits selective recognition of malignant cells even where the underlying protein is equally abundant on normal tissue — precisely the scenario that limited CD48-targeting agents and motivates renewed interest in conformation-selective formats. The atomic-resolution structural templates generated by crosslink-guided refinement provide a rational starting point for engineering such selectivity into antibodies, bispecifics, or cell-based therapies. As conformation-selective antibody discovery platforms mature, the pipeline established here — from experimental crosslinks to structural violation mapping to refined conformational models — could be applied systematically across the 212 candidate epitopes identified, prioritizing those with the sharpest conformational distinction between malignant and normal cell surfaces.

Structural surfaceomics addresses critical limitations in both traditional proteomics and structural biology approaches. The integration of two orthogonal crosslinkers (DSSO and PhoX) with complementary enrichment strategies provides both technical validation and expanded structural coverage, with consensus crosslinks showing lower violation rates (5.9%) compared to single-crosslinker observations (16.9%). The superior performance of high-pH reversed-phase fractionation likely reflects better separation of low-abundance crosslinked peptides based on hydrophobicity, and its offline stage-tip format is well-suited to low-input XL-MS samples. Importantly, proteins with mixed crosslinking distance patterns proved especially informative for integrative modeling and AI-enhanced tools. Our results highlight that current AlphaFold-derived integrative tools such as AlphaLink, although powerful on many targets, can fail catastrophically on a subset of proteins, particularly in the context of cell-surface conformations constrained by cell surface XL-MS. This discrepancy underscores that XL-MS-informed deep learning models should not be treated as ground truth; instead, they require orthogonal validation and careful diagnostic metrics at proteome scale. We therefore propose using structural surfaceomics not only for target discovery but also as a stress-test for emerging XL-MS-integrative algorithms, revealing regime boundaries where they fail to represent biologically validated conformational states.

While our approach identified numerous unique crosslinks exclusive to cancer cells, several limitations warrant consideration. The 30 Å distance constraint for crosslink validation, though standard, may miss subtle but functionally important conformational changes. Additionally, our analysis captures population-level conformational states rather than single-cell heterogeneity, potentially obscuring rare but therapeutically relevant conformations. Due to the low abundance of crosslinked peptides, another limitation of this method is the sample input requirements, which leads to variable coverage across the models. We note that many of the cancer-specific crosslinks could be a function of improved coverage in myeloma cell line models. Future iterations could incorporate recently developed crosslinkers with varied spacer lengths, single-cell structural proteomics approaches, and time-resolved measurements to capture conformational dynamics. Integration with cryo-EM structures and expanded computational modeling pipelines will further refine our understanding of these cancer-specific conformations. Going forward, to develop novel therapies and define new biology, advancements will be needed toward detection or analytical reagents that allow for rapid, orthogonal validation of potential alternative, tumor-specific conformations. Our mass spectrometry and structural modeling approaches presented here present an important milestone along this path.

The systematic identification of cancer-specific conformational epitopes opens new avenues for immunotherapy development. Unlike traditional immunotherapy targets that rely on differential expression, conformational epitopes offer the potential for truly tumor-selective recognition. The structural models generated through crosslink-guided refinement provide atomic-level templates for rational antibody design, potentially accelerating the development of conformation-selective therapeutics. Moreover, the database of 5,209 crosslinks across multiple cancer types serves as a foundational resource for the cancer structural biology community, enabling hypothesis generation and target prioritization. As protein structure prediction tools continue to improve, the integration of experimental crosslinking data will become increasingly valuable for identifying and validating conformational targets.

Ultimately, structural surfaceomics moves target discovery beyond protein expression to protein conformation, enabling a new class of precision immunotherapy where tumor selectivity is achieved not by differential abundance but by the malignancy-driven shapes proteins adopt at the cell surface.

## METHODS

### Cell culture and authentication

Leukemic suspension cell lines (AMO1, MM1.S, THP-1, MV411, NOMO-1) were cultured in RPMI-1640 supplemented with 10% FBS and 1% penicillin/streptomycin. Adherent prostate cell lines (PNT2, 22RV1) were cultured in glucose-dense RPMI-1640 (ATCC-modified) with 10% FBS and 1X penicillin/streptomycin. All cells were maintained at 37°C with 5% CO □, confirmed mycoplasma-free by MycoAlert (Lonza), and authenticated by STR profiling (12/05/2024, UC Berkeley DNA Sequencing Facility).

### Primary PBMC isolation

Peripheral blood mononuclear cells (PBMCs) were isolated from a healthy donor leukopak (StemCell Technologies) by density gradient centrifugation using Ficoll-Paque Plus (Cytiva/GE Healthcare). Leukopaks were diluted 1:5 with sterile phosphate-buffered saline (PBS) and carefully layered over Ficoll-Paque Plus in 50 mL conical tubes. Samples were centrifuged at 400 × *g* for 30 minutes at room temperature with the brake off. The buffy coat containing PBMCs was carefully collected from the plasma-Ficoll interface and washed twice with PBS by centrifugation at 300 × *g* for 10 minutes. Cell viability was confirmed to be >80% by trypan blue exclusion, and cells were used immediately for experiments.

### Live-cell surface crosslinking and biotinylation

Suspension cells were harvested at 80-90% confluency, washed three times with PBS, and resuspended in PBS at a concentration of 100 million cells per sample. For crosslinking, cells were treated with either 10 mM disuccinimidyl sulfoxide (DSSO, Sigma-Aldrich) or 10 mM PhoX (DSPP, TherThermo Fisher Scientific) as the final concentration, from a 50 mM stock solution resuspended in anhydrous dimethyl sulfoxide (DMSO, Sigma-Aldrich). Samples were incubated at room temperature for 45 minutes with rotation. Reactions were quenched with Tris-HCl pH 8.0 to a final concentration of 20 mM for 15 minutes, then washed twice with PBS (300 × *g*, 5 minutes).

Adherent cells were crosslinked in an on-plate format as to not disrupt the surfaceome by adding PBS washes and the crosslinking solution directly to the dish. After sample incubation and quenching the reaction, cells were gently detached from plates with a collagenolytic enzyme solution (Accutase, StemCell Technologies) to preserve cell surface epitopes, then washed three times with PBS, and transferred to microcentrifuge tubes for biotinylation.

For surface biotinylation using the biocytin hydrazide method, briefly crosslinked cells were resuspended in 990 µL cold PBS and treated with 1.6 mM sodium metaperiodate (NaIO □) for 20 minutes at 4°C with rotation to oxidize cell surface glycans as previously described^18,58^. Cells were washed twice with cold PBS and resuspended in 989 µL cold PBS containing 1 µL aniline, and samples were incubated with 1 mM biocytin hydrazide (Biotium) for 90 minutes at 4°C with rotation to covalently couple biotin to oxidized glycans on surface proteins. Cells were washed three times (500 × *g*, 3 minutes) before lysis and affinity purification.

For select DSSO-linked myeloma cell line samples (AMO1 and MM1.S), an alternative biotinylation method using WGA-HRP (Vector Laboratories) was tested. Following crosslinking and quenching as described above, cells (100 million per sample) were washed twice and resuspended in 500 µL pre-warmed PBS at 37°C. WGA-HRP was added to a final concentration of 0.04 mg/mL, followed by addition of biotin phenol (Sigma-Aldrich) to 0.5 mM final concentration. After gentle mixing, hydrogen peroxide (H□O□) was added to a final concentration of 10 mM to initiate peroxidase-catalyzed biotinylation. Samples were incubated at 37°C for 2 minutes with periodic gentle mixing. Reactions were quenched with ice-cold buffer (10 mM sodium ascorbate, 5 mM sodium pyruvate), and cells were washed with PBS three times before lysis or storage at −80°C.

### Biotinylation validation

Biotinylation efficiency was assessed by flow cytometry. Cells (1 × 10□) were stained with streptavidin-Alexa Fluor 647 (1:100, Invitrogen) for 30 minutes at 4°C, washed twice, and analyzed on a CytoFlex cytometer (Beckman Coulter). Data were processed using FlowJo (v10.9.0).

To verify specific enrichment of surface-biotinylated proteins and confirm successful crosslinking, lysates from biotinylated and crosslinked cells (PBMC-DSSO and MM1.S-DSSO) were prepared as described above and then subjected to immunoprecipitation with 10 µL monoclonal anti-CD48 antibody (clone MEM-102, Invitrogen) overnight at 4°C, followed by Protein A/G bead (Thermo Fisher Scientific) capture for 2 hours at 4°C. Beads were washed four times with RIPA buffer, and bound proteins were eluted by boiling in SDS-PAGE buffer, resolved by SDS-PAGE, transferred to PVDF membranes using the Turbo Blot system (Bio-Rad) for western blot detection with anti-CD48 (1:1000, Bethyl Laboratories) followed by anti-rabbit HRP-conjugated secondary antibody (diluted 1:20, Cell Signaling Technologies). Bands were detected via ECL and the blot was imaged (ChemiDoc Imaging System).

### Streptavidin affinity enrichment

Biotinylated cell pellets were lysed in 1 mL of 2× RIPA buffer (20 mM Tris-HCl pH 7.5, 300 mM NaCl, 2% NP-40, 1% sodium deoxycholate, 0.2% SDS) with 1× HALT protease inhibitors (Thermo Fisher Scientific) and 2 mM EDTA. Lysates were sonicated on ice until clarified (20-second pulses, 1-2 minutes total), incubated 10 minutes on ice, then centrifuged (17,000 × *g*, 10 minutes, 4°C).

For affinity enrichment, 650 µL of High Capacity NeutrAvidin agarose resin slurry (Thermo Fisher Scientific) per sample was transferred to 10 mL gravity flow columns using wide-bore pipette tips. Resin was washed three times with 1 mL of 1× RIPA buffer containing 1 mM EDTA. Then, cell lysates were added to the column and incubated with end-over-end rotation for 2 hours at 4°C. Following incubation, resin was washed sequentially by gravity flow with approximately 50 mL each of: (1) 1× RIPA buffer containing 1 mM EDTA, (2) PBS containing 1 M NaCl to disrupt non-specific interactions, and (3) 50 mM ammonium bicarbonate (ABC) containing 2 M urea to remove residual detergents. After final washing, resin was prepared for on-bead proteolytic digestion.

### Proteolytic digestion and mass spec preparation

Washed resin was treated with 250 µL one-pot proteolytic digestion buffer (50 mM Tris pH 8.5, 10 mM TCEP, 20 mM IAA (iodoacetamide), 4 M urea, pH verified to 8.0-8.5) and incubated with 25 µg trypsin/Lys-C (Mass Spec Grade, Promega) at 37°C for 2 hours. After diluting to 1 M urea with 50 mM Tris-HCl buffer, digestion continued overnight at room temperature. Peptides were collected, beads were rinsed with 500 µL ABC, then solution was acidified with 1% TFA to pH <3, and centrifuged (17,000 × *g*, 5 minutes). Peptides were desalted using SOLA HRP cartridges (Thermo Fisher Scientific) conditioned with ACN and 0.1% TFA, washed with 0.1% TFA and 2% ACN/0.1% FA, then eluted with 80% ACN/0.1% TFA. Eluates were dried in a SpeedVac and stored at −80°C. until further fractionation or LC-MS/MS analysis.

### Crosslinker-specific enrichment

#### IMAC (PhoX samples)

Dried peptides were resuspended in 200 µL binding buffer (80% ACN, 0.1% TFA) and loaded onto Fe³□-NTA spin columns (Thermo Fisher Scientific). After 30-minute incubation with gentle mixing, columns were washed three times with binding buffer and once with MS-grade water. Peptides were eluted twice with 100 µL ammonium hydroxide buffer (5%), dried by vacuum centrifugation, and stored at −80°C.

#### SEC (DSSO samples)

Peptides were reconstituted in 25 µL SEC buffer (30% ACN/0.1% TFA) and separated on a Superdex Peptide 3.2/300 column (Cytiva) at 50 µL/min for 90 minutes using an Agilent HPLC. Fractions were collected every 2 minutes (45 fractions total). Fractions 14-15 containing crosslinked peptides were pooled, dried, and stored at −80°C.

### In-stage tip high pH reverse phase fractionation

To reduce sample complexity prior to LC-MS/MS analysis, enriched crosslinked peptides were fractionated by high pH reversed-phase chromatography using stage-tip methodology. Stage-tips were prepared by packing approximately 5 mg of high pH compatible C18 beads (Durashell 150 A, 3 μm, Phenomenex) over a C18 solid phase extraction disk (Empore) into 200 µL pipette tips, which were then conditioned by sequential addition of 90 µL methanol and 150 µL ACN (partially drained) via centrifugation at 1,200 × g for 4-5 minutes. Tips were filled with ACN and equilibrated for at least 2 hours, then washed with pH 10 buffer (25 mM NH OH). Peptides dissolved in pH 10 buffer were loaded, washed once, and eluted with increasing ACN (6-50% in pH 10 buffer). Fractions were concatenated (25%+6%, 30%+9%, 35%+12%, 50%+21%) to yield six fractions, dried, and stored at −80°C.

### LC-MS/MS data acquisition

**TimsTOF Pro 2 Analysis** Crosslinked peptide fractions were analyzed on a TimsTOF Pro 2 mass spectrometer (Bruker Daltonics) coupled to a nanoElute UHPLC system. Chromatographic separation utilized a 25 cm × 75 μm PepSep C18 column (1.6 μm particles) maintained at 50°C. A binary gradient of 0.1% formic acid in water (A) and 0.1% formic acid in acetonitrile (B) was employed over 30 minutes at 400 nL/min: 2-27% B (0-100 min), 27-45% B (100-110 min), 45-95% B (110-115 min), followed by column wash and re-equilibration. The TIMS device operated with 1/K_0_ scan range from 0.70 to 1.30 Vs/cm², 100 ms ramp time, and PASEF acquisition mode. MS1 spectra were acquired from 100-1700 *m/z* with ion mobility resolution optimized for crosslinked peptide separation. Data-dependent PASEF selected precursors with charges 3-8, applying dynamic exclusion for 0.4 minutes. CID fragmentation was performed with collision energy ramped linearly based on ion mobility.

#### Orbitrap Astral High-Resolution Analysis

High-sensitivity measurements of crosslinked peptide fractions were performed on an Orbitrap Astral mass spectrometer interfaced with a Vanquish Neo UHPLC. Peptides were resolved on a 25 cm × 75 μm PepMap C18 column (2 μm particles) at 40°C using a 50-minute gradient of 0.1% formic acid in water (A) and 0.1% formic acid in 80% acetonitrile (B) as follows: 4-22.5% B over 37.5 minutes, then 22.5-35% B over 12.5 minutes at 250 nL/min. Survey scans covered 400-1400 *m/z* at 180,000 resolution (FWHM) with 500% normalized AGC and 5 ms maximum fill time. Data-dependent MS2 acquisition operated in 1-second cycles, targeting charge states ≥3 with 1.6 *m/z* isolation width. Fragment ion spectra (150-2000 *m/z*) were acquired at 80,000 resolution using 32% normalized HCD energy, 500% AGC target, and 20 ms maximum injection time. A 15-second dynamic exclusion window prevented redundant sampling.

#### Orbitrap Eclipse DIA for surfaceomics (non-XL)

Un-crosslinked surfaceomics samples were analyzed via data-independent acquisition on an Orbitrap Eclipse Tribrid system coupled to a Vanquish Neo UHPLC with a 60 cm x 75 μm IonOpticks C18 column (2 um particles, Aurora series) operated at 300 nL/min and 1500 bar maximum pressure. The 135-minute method employed sequential gradient elution: 3-25% B (0-102 min), 25-40% B (102-122 min), 40-90% B (122-123 min), maintaining 90% B (123-125 min). Precursor scans (380-985 *m/z*, 60,000 resolution) preceded 51 consecutive DIA windows spanning 300-1200 *m/z* with 12 *m/z* isolation width and 1 m/z overlap. MS2 spectra were acquired at 30,000 resolution using 25% HCD energy, automatic AGC, and 40 ms maximum injection.

### Crosslink identification and validation

#### XL-MS data

Raw files were searched using pLink 2.3.11 for crosslink identification. Search parameters included: human UniProt database (2024_01 release), trypsin specificity allowing 3 missed cleavages, carbamidomethylation as fixed modification, and methionine oxidation/N-terminal acetylation as variable modifications. Crosslinker-specific modifications were defined for DSSO (158.0038 Da) and PhoX (209.97181 Da) with appropriate cleavable sites. Mass tolerances were set to 10 ppm (precursor) and 20 ppm (fragment) for Orbitrap instruments, with adaptive tolerances for TimsTOF data. FDR thresholds of 5% were enforced at the peptide pair level using target-decoy approaches.

#### Filtering and dataset statistics

After filtering for common contaminants and low-quality spectra, the redundant crosslink spectral matches (CSMs) were aggregated to unique crosslinks across datasets for inter-dataset quality control, reproducibility metrics, and dataset-level statistics to ensure reproducibility and minimize false discoveries. CSMs from each dataset were filtered at 1–5% FDR using pLink prior to combining, and unique crosslinks were defined by unique protein pair + residue position combinations across all 11 datasets. Inter-replicate reproducibility was assessed by Jaccard similarities, which ranged from 0.18-0.87, with myeloma samples demonstrating the strongest concordance (mean similarity = 0.50).

#### Label free quantification

For paired shotgun surfaceomics (without crosslinking) before and after streptavidin purification, DIA data were processed with DIA-NN 1.8.1 (library-free mode). DDA data were analyzed with FragPipe 22.1 and MaxQuant 8.0 using standard LFQ workflows with match-between-runs enabled to generate protein ranks, confirming significant enrichment of membrane proteins (Gene Ontology cellular component analysis, p<0.001).

### Disease-specific crosslink classifications

We implemented a multi-tier validation strategy including overlap and principal component analysis, dendrogram clustering with z-score calculation. Disease-specific crosslinks were defined using a stringent two-tier criterion: (1) detection across cancer models, and (2) absence from other models. Crosslinks were also classified by crosslinker consensus validation. Protein networks were analyzed using STRING, with statistical enrichment via KEGG and GO analyses.

### Structural analysis

#### Distance calculations

The Euclidean Cα-Cα distances for each crosslink were calculated using AlphaFold2 structures with our tool AlphaCross-XL^22^. Crosslinks were classified as satisfied (≤30 Å) or violated (>30 Å) and visualized in PyMOL along with the protein pLDDT score (v2.5, Schrödinger). Predicted aligned error (PAE) matrices from AlphaFold predictions were visualized using the PAE Viewer^59^ to assess domain-domain confidence and identify flexible regions.

#### Computational environment

All conformational refinement was performed on an Amazon Web Services (AWS) g5.2xlarge instance (8 vCPUs, 32 GB RAM) equipped with an NVIDIA A10G GPU (24 GB VRAM). GPU support used NVIDIA driver 575.51.03 with CUDA 12.9 (CUDA compiler tools v12.9.41). All analyses were executed in a conda-managed environment (openfold-new) to ensure reproducible dependencies.

#### Conformational refinement

Proteins with disease-specific crosslink patterns inconsistent with AlphaFold structures were refined using AlphaLink^33^ (via OpenFold^60^) and MODELLER v10.7 (Šali Lab, UCSF)^27,52,53^. AlphaFold2-predicted structures served as starting templates for comparative modeling with custom spatial restraints derived from experimental crosslinking data. For CD48, flexible regions (residues 114-117, 128-135) were refined under experimental distance restraints (Lys98-Lys114 ≤25 Å; residues 128-135 closed conformation). Twenty-five models were generated per region, ranked by DOPE score, validated with MolProbity, and agreement with XL-MS data.

#### Post-prediction relaxation

To resolve steric clashes and improve stereochemical quality, predicted structures were energy-minimized using OpenMM (v8.4) via OpenFold relaxation utilities. Missing atoms and hydrogens were added using PDBFixer at physiological pH (pH 7.0) prior to minimization. Energy minimization was performed with positional restraints applied to non-hydrogen atoms (restraint set “non_hydrogen”), using a restraint stiffness of 10.0 kcal mol^-1^ Å^-2^ and an energy/length tolerance of 2.39 kJ mol^-1^ nm^-1^. Minimization was executed using the OpenMM CPU platform for consistent execution in the deployed environment. Relaxed structures were written in PDB format while preserving residue numbering and pLDDT values.

#### Crosslink network analysis and visualization

To assess the biological context and confidence of detected crosslinks, protein-protein interaction networks were constructed using XiNet^61^ with STRING database annotations, and crosslink patterns were overlaid on known complexes. XL-Ranker^62^ was employed to resolve peptide-mapping ambiguity.

#### Protein-protein complex modeling

For inter-protein crosslinks, protein-protein complexes were modeled using multiple complementary approaches. AlphaFold-Multimer^48^ was used to predict structures of protein complexes as starting templates, which were then compared with enhanced predictions using AlphaLink2^23^ (based on Uni-Fold Multimer^63^), which incorporates crosslink distance restraints directly into the structure prediction workflow. Models were evaluated by PAE and ipTM scores.

#### Comparative modeling of integrin VLA-4

We computed a comparative model of the integrin VLA-4 (α4/β1) heterodimer in the open conformation using MODELLER^52,53^. Templates were identified using PDB BLAST and HHpred searches on the Protein Data Bank (PDB). The model used multiple templates, including the α and β chains of the cryoEM structure of integrin α4/β7 (PDB code 9P95), the α chain of the cryoEM structure of integrin α5/β3 (PDB code 9JEI), the β chains of the cryoEM structure of integrin α5/β1 (PDB code 7NWL), and the X-ray structure of the extended integrin β3 (PDB code 6BXB). Sequence-structure alignments were constructed with PROMALS3D^64^.

## DATA AVAILABILITY

The raw files of the proteomic data along with the supporting processed data generated for this article here have been deposited at the ProteomeXchange/PRIDE repository with accession numbers PXD059495, PXD035589, and PXD035591. The final mass spectrometry datasets are pending due to temporary repository service interruption and the working submission reference is 1-20260506-005651-1000006434.

## CODE AVAILABILITY

The code generated for this article here is available at https://github.com/audrey-kishishita/structural_surfaceomics.

## ACKNOWLEDGMENTS

We thank members of the laboratory of A.P.W. for feedback and support; C.K. for routine mycoplasma testing of cell lines; A.I. for routine cell line authentication and UC Berkeley DNA Sequencing Facility for STR profiling; and S.R. for oversight of CytoFLEX operations and cryogenic storage systems. We thank members of the laboratory of B.Z. for LC-MS operation and instrument support (timsTOF Pro2). We are grateful to the instrumentation managers in the UCSF Antibiome Center, including S.K. for HPLC system support, and T.P.C., K.L., and S.T. for LC-MS system management (timsTOF Pro2). We thank the field service engineers at Thermo Fisher Scientific for technical support on LC-MS/MS instrumentation (Vanquish Neo, Orbitrap Eclipse). We also thank Tom Goddard of UCSF RCVI and the UCSF Wynton team for suggestions, feedback and GPU-enabled computational support. This work was supported by NIH grant 1R01CA290875-01 (to A.P.W. and L.H.).

## AUTHOR CONTRIBUTIONS

A.K. conceived the study and designed the overall research strategy together with A.P.W. and K.M. A.K. performed XL-MS sample preparation, investigation, data acquisition, formal analyses, and figure generation with assistance from S.C. T.G. performed dataset-wide AlphaLink processing. S.P. performed figure visualization work, western blot optimization, and figure enhancement. R.D. carried out peptide localization and structural enrichment analyses. C.Y., I.R., S. Shenoy, A.Z., M.R.H., D.B., and R.L.M. contributed computational analyses and software support. A.B., A.R., H.L., and A.N. generated proteomics datasets used in the abundance rank plot. F.J. assisted with high-pH fractionation and MS3 method development. Y.H. performed sample acquisition on the mass spectrometer and contributed formal data analysis. M.H. and C.S. carried out structural modeling of Integrin complexes. R.D. provided HPC cluster support. S.B. contributed flow cytometry experiments and analysis. F.F., L.H., S. Srivastava, A. Sali, R.V., K.M., and A.P.W. supervised various aspects of the study and provided critical resources. A.K. and A.P.W. wrote the manuscript with input from all authors.

## CORRESPONDING AUTHOR

Correspondence to Arun P. Wiita, MD, PhD.

