## Supplemental Information for "Extending structural surfaceomics to identify aberrant conformations of tumor surface proteins as potential immunotherapy targets"

Audrey Kishishita et al.

**Supplementary Figures**

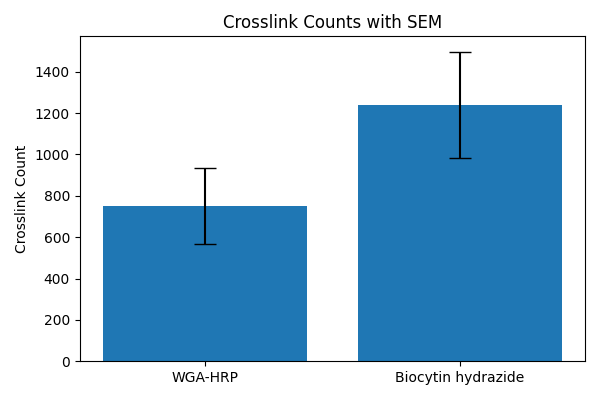

**Supplementary Figure 1. Comparison of biotinylation methods for DSSO-crosslinked blood cancer suspension cells.**

Bar plot shows mean unique crosslink counts (±SEM, n=2 biological replicates) for WGA-HRP proximity labeling on DSSO-myeloma lines versus biocytin hydrazide glycan oxidation approaches. Biocytin hydrazide yielded 1.65-fold more crosslinks (1,240 ± 257) compared to WGA-HRP (750 ± 183), demonstrating superior surface protein enrichment efficiency for structural surfaceomics workflows.

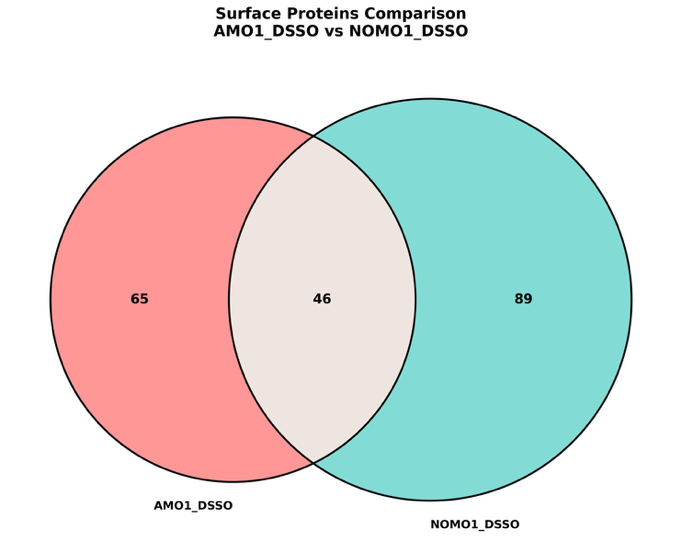

**Supplementary Figure 2. Biocytin hydrazide and WGA-HRP biotinylation methods identify similar surface protein complements.**

Venn diagram comparing surface protein identification from DSSO-crosslinked cells using WGA-HRP proximity labeling (AMO1, red) versus biocytin hydrazide N-glycan oxidation (NOMO-1, teal). While both methods revealed similar numbers of GO identified surface proteins (111 vs 135 proteins, respectively), with 46 proteins shared between approaches, biocytin hydrazide yielded substantially more crosslinks per protein (Supplementary Fig. 1), demonstrating superior depth of structural coverage despite comparable surfaceome.

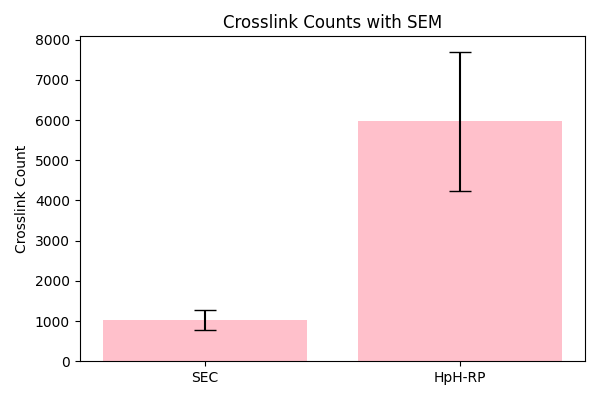

**Supplementary Figure 3. High-pH reversed-phase (HpH-RP) fractionation enhances PhoX crosslink detection compared to size exclusion chromatography (SEC).**

Evaluation of second-dimension fractionation strategies following Immobilized Metal Affinity Chromatography (IMAC) enrichment of PhoX-crosslinked peptides. Bar plot shows mean unique crosslink counts (±SEM, n=2 biological replicates) from NOMO-1 (AML) cells processed by SEC versus myeloma cells processed by HpH-RP (high-pH reversed-phase) chromatography. HpH-RP fractionation yielded 5.8-fold more total crosslinks (5,965 ± 1,733) relative to SEC (1,028 ± 250), establishing IMAC-HpH-RP as the optimal workflow for PhoX-based structural surfaceomics.

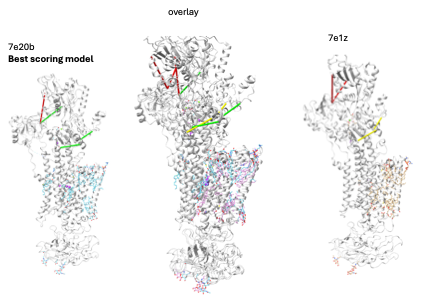

**K^+^ E2 state**

**Na^+^ E1 state**

**Supplementary Figure 4.** For proteins with multiple experimental structures, we applied xlms-tools^2^ to identify structures matching our crosslink data. In the example above of Sodium Na^2+^/K^+^ pump, the PBD: 7e20b is the best scoring model.

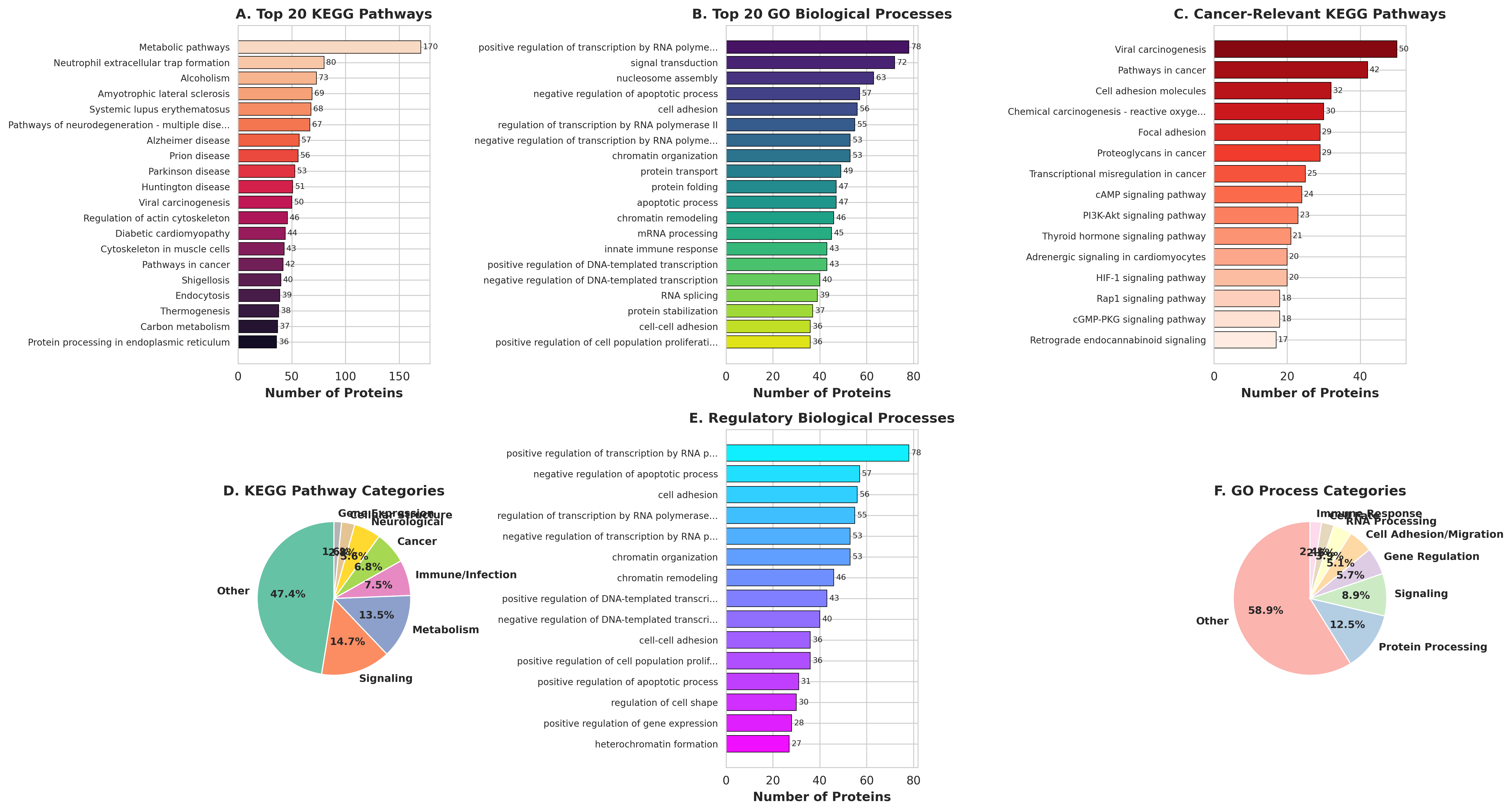

**Supplementary Figure 5.** Functional Enrichment Analysis of Proteins Harboring Distance-Violating Crosslinks. Pathway and gene ontology (GO) analyses were performed on proteins with at least one high-confidence distance-violating crosslink (pLDDT > 70) to identify biological processes and pathways associated with cancer-specific conformational remodeling. **(A)** Top 20 KEGG pathways by protein count. Metabolic pathways represent the largest category (n = 179), followed by neutrophil extracellular trap formation (n = 99) and neurodegeneration-associated pathways (Alzheimer, Parkinson, Huntington disease). Cancer-relevant pathways, including viral carcinogenesis and pathways in cancer, appear among the top 20, alongside cell adhesion and cytoskeletal regulation. **(B)** Top 20 GO Biological Processes. Transcriptional regulation dominates (positive regulation of transcription by RNA polymerase II, n = 78; signal transduction, n = 72), followed by nucleosome assembly, negative regulation of apoptotic process, and cell adhesion (n = 58–63). Chromatin organization, remodeling, and protein transport are also prominently represented, indicating broad involvement of gene-regulatory and structural processes. **(C)** Cancer-relevant KEGG pathways subset. Viral carcinogenesis (n = 59) and pathways in cancer (n = 42) are the most enriched. Cell adhesion molecules (n = 37), focal adhesion (n = 29), proteoglycans in cancer (n = 29), and PI3K-Akt signaling (n = 23) further highlight cancer-relevant functional categories, consistent with roles for conformationally remodeled proteins in tumor adhesion and oncogenic signaling. **(D)** Categorical breakdown of all enriched KEGG pathways. Signaling (14.7%) and metabolism (13.5%) are the largest defined categories after the broad "other" group (47.4%), with immune/infection (7.5%) and cancer pathways (6.8%) also well-represented. **(E)** Regulatory GO biological processes subset, highlighting chromatin remodeling, transcriptional regulation, apoptosis regulation, and cell adhesion as the predominant regulatory categories among violating proteins. **(F)** Categorical breakdown of enriched GO biological processes. Protein processing (12.5%) and signaling (6.9%) represent the largest functional categories after "other" (58.9%), with cell adhesion/migration (5.7%) and gene regulation also enriched, consistent with the pathway-level findings in panels A–C.

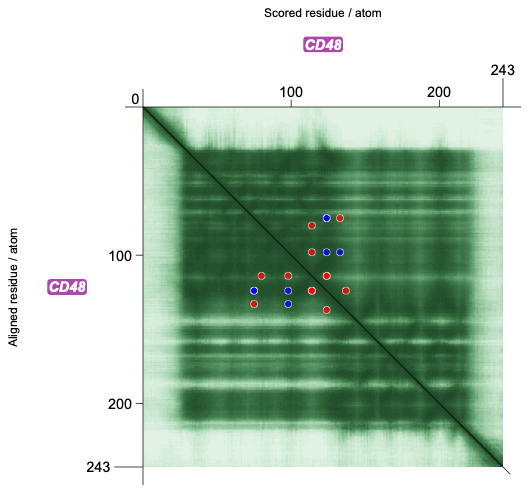

**Supplementary Figure 6. Predicted aligned error (PAE) analysis of CD48 with mapped crosslinks.** PAE plot for CD48 (AlphaFold2 prediction) showing crosslink positions overlaid on the confidence matrix. Each dot represents a detected crosslink, color-coded by distance constraint validation: blue dots indicate satisfied crosslinks (Cα-Cα distance ≤30 Å), red dots indicate violated crosslinks (>30 Å). The diagonal line represents self-alignment (dark green, zero error). Darker green regions indicate higher confidence in relative domain positioning, while lighter regions suggest greater structural uncertainty. Distance-violating crosslinks (red) map predominantly to high-confidence structural regions with variability (medium green), supporting the interpretation that violations reflect genuine conformational differences rather than AlphaFold prediction uncertainty.

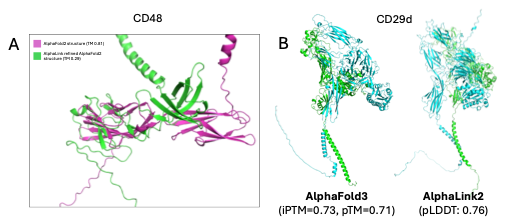

**Supplementary Figure 7. Conformation-specific targets reveal failure modes of AlphaLink.
(A)** For the candidate protein CD48, overlay of the AlphaFold2 model (*magenta*) with the AlphaLink‑refined model (*green*) shows pronounced loss of secondary structure and distortion of the folded domains (*black arrows*). Relative to the crystal structure homolog (PDB: 2DRU), the AlphaLink model exhibits a substantially reduced TM‑score (0.29) compared to the original AlphaFold2 model (0.81), indicating degradation of global topology rather than refinement.
**(B)** For the candidate protein complex CD29d, the AlphaFold3 model (*left*) displays well‑folded domains with acceptable iPTM and pTM scores (0.73 and 0.71, respectively). After AlphaLink2 optimization, the model retains an overall acceptable pLDDT score (0.76) but shows extensive misfolding and domain distortion, indicating that the refinement failed to capture the hypothesized conformational dynamics and instead generated structurally implausible solutions.

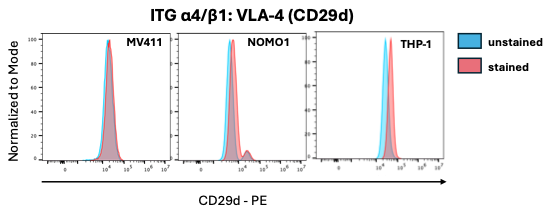

**Supplementary Figure 8. Flow cytometric detection of active VLA-4 conformation on AML cell lines.** Surface expression of active Integrin α4/β1 (VLA-4) was assessed using the CD29d antibody (PE-conjugated), which selectively recognizes the extended, activation-associated epitope on the β1 integrin subunit (ITGB1)^2^. Representative histograms are shown for three AML cell lines: MV4-11, NOMO1, and THP-1. Unstained cells (blue) served as a reference for background fluorescence. Stained cells (pink) are shown overlaid. THP-1 cells exhibit a clear rightward shift relative to unstained controls, indicating robust surface expression of the active β1 conformation. MV4-11 and NOMO1 show lower but detectable staining above background. These data support the presence of the open, extended VLA-4 headpiece conformation on AML cell surfaces, consistent with inter- and intra-subunit crosslinks that violate the AF3 closed-conformation model.
